# Sustained epithelial interferon signaling modulates incomplete pathologic response in colorectal cancer

**DOI:** 10.1101/2025.03.24.644737

**Authors:** Erdogan Pekcan Erkan, Emmi Hämäläinen, Julia Kolikova, Kornelia Kuc, Kalle Ojala, Meiju Kukkonen, Ismail Hermelo, Selja Koskensalo, Timo Tarvainen, Alli Leppä, Ilona Keränen, Carola Haapamäki, Essi Karjalainen, Monica Carpelan-Holmström, Laura Renkonen-Sinisalo, Laura Koskenvuo, Pauli Puolakkainen, Jukka-Pekka Mecklin, Elina Virolainen, iCAN, Matti Nykter, Lauri A. Aaltonen, Anna Lepistö, Ari Ristimäki, Toni T. Seppälä

**Author notes:** Correspondence: Toni Seppälä, MD PhD., Faculty of Medicine and Health Technology, Tampere University, Arvo Ylpön katu 34, 33520 Tampere, Finland. Co-first authors – equal contribution (Erdogan Pekcan Erkan, Emmi Hämäläinen).

## Abstract

**Background & Aims:** Patients with colorectal cancer have heterogeneous clinical responses to chemotherapy, although clinical guidelines advise little variability in treatment selection based on molecular tumor features. Precision oncology research typically utilizes patient-derived tumor organoids (PDTO) to predict clinical outcomes, but such efforts are often not directed towards identification of molecular factors underlying differential responses to therapy.

**Methods:** Bulk RNA-sequencing was performed on treatment-naive PDTOs, and gene expression data was combined to drug sensitivity data to identify transcriptomic features associated with low *in vitro* sensitivity to chemotherapy. Whole-exome sequencing was performed on primary tumors to infer the somatic mutations of PDTOs and used to identify somatic mutations associated with differential *in vitro* drug responses. Publicly available gene expression and drug sensitivity data sets were used to validate the results. RNA interference was used for functional validation.

**Results:** PDTOs with low chemosensitivity had high JAK-STAT pathway activity resulting from high expression of interferon-stimulated genes. Evidence from single-cell RNA-sequencing confirmed chemotherapy-induced expression of interferon-stimulated genes in epithelial cells of cancers with partial response. *EPSTI1* knockdown decreased cancer cell viability and sensitized cells to chemotherapy.

**Conclusions:** Sustained interferon signaling in epithelial cancer cells contributes to incomplete pathologic response in colorectal cancer. The findings highlight the potential of JAK-STAT inhibition or TRAIL pathway activation to enhance chemotherapy efficacy. Future studies investigating pharmacologic modulation of these pathways in preclinical CRC models are needed to determine their viability as therapeutic targets.

## INTRODUCTION

Patients with colorectal cancer (CRC) can have heterogeneous clinical responses to conventional chemotherapeutics, although clinical guidelines advise little variability in treatment selection based on molecular tumor features. Primarily composed of epithelial cells^1^, patient-derived tumor organoids (PDTOs) have emerged as preclinical models that are particularly suited for studying epithelial cell-specific features, since they accurately recapitulate the histopathological and genomic characteristics of tumor tissues they are derived from^2–4^. In functional precision oncology, PDTOs have been used to study tumor progression and metastasis^5^, and to evaluate and predict patient-specific responses to chemotherapy^6,7^ and immunotherapies^8^. Yet, these models have been underutilized to identify molecular features of epithelial cancer cells and understand how they respond to chemotherapeutics.

Here, we performed integrative molecular profiling of primary CRC tumors and matched PDTOs to identify genomic and transcriptomic factors underlying heterogeneous *in vitro* drug responses to conventional chemotherapy. Our findings link epithelial interferon signaling to low chemosensitivity to study the determinants of incomplete pathologic responses.

## MATERIALS AND METHODS

### Ethical issues

The study was approved by the ethical committee of the Helsinki and Uusimaa hospital district and study site institutional review boards (PANORG ethical approval HUS/390/2021, HUS approval HUS/69/2022). SYNCOPE trial was approved by the Helsinki and Uusimaa ethical committee, study site-specific institutional review boards and national Finnish Medicines Agency FIMEA (ethical approval HUS/2427/2020 and HUS approval HUS/155/2021, trial registration NCT04842006, EudraCT 2020-003697-52, CTIS transition EU CT 2024-517149-15-00 and Fimea/2024/005474). The study participants provided a written, informed consent for the protocols.

The iCAN Flagship Project biobank study is based on a biobank consent and was reviewed by the HUS Ethical Committee and is executed based on a HUS research permit (4.5.2023 §38 (HUS/223/2023), a Findata data permit (THL/1338/14.02.00/2022), and MTAs with Helsinki Biobank (HBP20210170) and Finnish Hematology and Registry Biobank FHRB (12.5.2022). The project is carried out entirely in the HUS Acamedic safe processing environment where the patients remained non-identifiable.

### Patient recruitment

This study was carried out as part of PDTO tissue acquisition protocols: PANORG project and SYNCOPE clinical trial at Helsinki University Hospital. Patients enrolled on PANORG protocol also participated in the iCAN Flagship Project through collection of biobank samples and biobank transfer of PANORG samples.

Patients were enrolled to the study protocols at their first outpatient visit to tertiary cancer care at Meilahti Hospital (Helsinki, Finland) or Jorvi Hospital (Espoo, Finland). Both hospitals are part of the HUS Comprehensive Cancer Center. Upon written informed consent, CRC tumor tissues and germline DNA reference were collected, with the primary function of molecularly characterizing primary tumors and establishing PDTO cultures.

### Tissue collection

Primary tumor tissue samples were obtained by a study surgeon either via endoscopic biopsy forceps or conchotome before any treatment, or via biopsy from surgical specimen potentially exposed to neoadjuvant therapy. The samples were transported to the laboratory site in a transfer medium containing DMEM supplemented with 1% Penicillin-Streptomycin solution to establish PDTOs. Tissue specimens were processed within 24 hours after collection to minimize the risk of autolysis and tissue degradation.

Primary CRC samples for exome and transcriptome sequencing were collected to Helsinki Biobank from the surgical resection specimen at the Department of Pathology. Peripheral blood mononuclear cell reference DNA samples were collected through biobank as part of the routine laboratory work-up for the participants.

### PDTO culture

Tumor tissue samples were dissociated mechanically with a P1000 pipette, followed by a ten-minute enzymatic digestion with Organoid Digestion Medium. Only mechanical dissociation was used for endoscopic biopsies. Dissociated samples were centrifuged and washed three times with Organoid Wash Medium. After the final centrifugation step, cell pellets were resuspended in Matrigel (Corning, NY, USA) and plated on a 24-well plate. The plate was incubated at 37 °C for 30 minutes, and 500 µL of Organoid Culture Medium supplemented with 1 µL/mL Rho-associated, coiled-coil containing protein kinase (ROCK) inhibitor Y-27632 was added. Our success rate to establish PDTOs from surgical specimens was on par with earlier reports of similar sample sizes^7,9,10^.

PDTOs were incubated at 37 °C with 21% O_2_ and 5% CO_2_ and subcultured every 7-10 days for expansion. The Wnt-activating niche factors Wnt-3a and R-spondin were not added to the Organoid Culture medium to select and support the growth of Wnt-independent tumor cells. The critical biomass for a stable organoid line for molecular and phenotypic characterization was achieved early in most cases, but the expansion phase was continued for 4-6 passages to ensure independence from Wnt-activating supplements. *In vitro* drug testing was not performed before passage 5 (P5) to minimize the risk of assaying non-cancerous PDTOs.

### Single-cell RNA sequencing (scRNA-seq) of PDTOs

Chromium Fixed RNA Profiling for Multiplexed Samples (10x Genomics) was used to perform single cell RNA sequencing (scRNA-seq) on PDTOs. Briefly, PDTOs were dissociated to single cells with TrypLE Express (12605036, Gibco) and filtered through a strainer. Chromium Next GEM Single Cell Fixed RNA Sample Preparation Kit (PN-1000414, 10x Genomics) was used to fix cells at 4 °C for 24 hours. Sequencing libraries were prepared by using the multiplexed workflow of the Chromium Fixed RNA Kit, Human Transcriptome (PN-1000476, 10x Genomics) sequenced on a NovaSeq 6000 instrument (Illumina) in paired-end mode.

The cellranger multi pipeline v.9.0.0 (10x Genomics) was used for demultiplexing, alignment, and counting. Seurat v.5.0 R package was used to create Seurat objects from the output files. In the first quality-control step, the miQC framework was used to filter out low-quality cells in a data-driven way. In the second quality-control step, cells 1) expressing less than 2500 features and 2) those expressing combinations of *EPCAM*, *FN1*, and *PTPRC* were further filtered out.

Downstream analyses were performed without integration, since PDTO samples were processed and sequenced at the same time. NormalizeData function was used to normalize raw counts (normalization.method = “LogNormalize”, scale.factor = 10,000). FindVariableFeatures function was used to identify the variable features, and ScaleData function was used to scale the normalized data. RunPCA function was used to perform principal component analysis (PCA).

FindNeighbors function was used for shared nearest-neighbor (SNN) graph construction (reduction = "pca”, k.param = 30, dims = 1:10) and FindClusters function was used to identify the clusters on the SNN graph across multiple resolutions. RunUMAP function was used to create a uniform manifold approximation and projection (UMAP) embedding.

### DNA isolation from primary tumors and whole exome sequencing

QIASymphony DSP DNA kit was used to isolate genomic DNA from tumor tissue samples and buffy coat of blood samples. Twist Exome 2.0 plus (Twist Exome 2.0 plus Comp Exome spike-in) was used for library preparation (input = 50 ng genomic DNA). Indexed libraries were pooled and paired end sequenced on an Illumina NovaSeq6000 system (Illumina, San Diego, CA, USA) (read length: 2 x 101 bp). Depth of sequencing was 30x for germline reference DNA (>90% of target covered with >30x) and 100x for tumor DNA (>95% of target covered with >100x).

### Secondary analysis (preprocessing) of whole exome sequencing data

The sequenced data was processed at Institute of Molecular Medicine Finland (FIMM) with Dragen DNA pipeline v3.10.8 (DRAGEN Host Software Version 4.0.3 and Bio-IT Processor Version 0x18101306) using GRCh38.105 human reference genome for alignment. The analysis was performed for germline only (using normal sample) and for somatic (using tumor-normal pairs).

### Exome variant categorization

DRAGEN DNA pipeline output ‘hard-filtered.vcf.gz’ files containing single-nucleotide variants and indels were used for annotating all somatic variants with VEP^16^, keeping only FILTER column ‘PASS’ variants. The vc2maf tool^17^ was used to convert the annotated *.vcf files to *.maf files, and the maftools R package^18^ was used to visualize the variants.

### *In vitro* drug testing

5-fluorouracil (S1209, Selleckchem), oxaliplatin (S1224, Selleckchem) and irinotecan (S2217, Selleckchem) were resuspended and stored according to manufacturer’s instructions. PDTOs were dissociated into single cells, resuspended in Organoid Culture Medium:Matrigel (9:1) mixture, and plated in a 384-well plate. Cells were let to recover for 48 hours, and organoid formation was visually assessed. Drugs were applied with a digital liquid dispenser (Tecan D300e,

Tecan, Zurich, Switzerland) across a logarithmically designed dose range spanning ten concentrations. Dimethyl sulfoxide (DMSO, max. 1%) was used as negative control for 5-FU and irinotecan, while water + 0.3% Tween-20 was used as negative control for oxaliplatin. Blasticidin (ant-bl-005, Invivogen) was used as a positive control at a final concentration of 10 µg/ml. Four technical replicates were used to correct for the intraplate variability. PDTOs were treated with drugs for five days, and CellTiter-Glo assay (Promega, Madison, WI, US) was used to assess cell viability on day 7. Luminescence signals were measured on a multimode plate reader (FLUOstar Omega, BMG Labtech).

### Drug response modeling

The average luminescence signal in the blank wells was subtracted from each well to calculate blank-corrected signals. Relative cell viability was calculated with the formula:

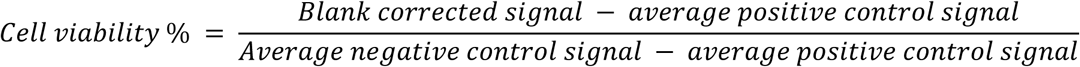

The four-parameter logistic model is arguably the most common statistical method to identify drug efficacy *in vitro*^11,12^ and it is often used regardless of the data satisfying the assumptions underlying the model, producing potentially inaccurate point estimates (IC50 or AUC) that are used for downstream analyses. Moreover, IC50 values can only be calculated if the relative cell viability is less than 50% at the highest drug concentration tested^13^, a requirement that was not met in many PDTOs in our study. Therefore, we used the nonparametric monotone (npM) model of the ENDS tool^11^ by fitting the model over the mean responses at each dose. AUC values were used as point estimates of *in vitro* drug responses.

### Quality control and drug response classification

Dose-response curves were visually evaluated. An assay was considered to have failed if the normalized cell viability was less than 50% at the lowest drug concentration or the nonparametric model failed (e.g., a flat line was seen). Twelve PDTOs had failed assays for all three drugs and were excluded from downstream analyses. Kernel density estimation (KDE) was used on AUC values to find two valley points to be used as thresholds for drug response classification. This approach found two valley points for fluorouracil, but not irinotecan and oxaliplatin. As the secondary approach, we used quartile-based classification for irinotecan and oxaliplatin:

**if** AUC > Q3 ∼ “low sensitivity”

**elif** AUC ≤ Q1 ∼ “high sensitivity”

**else ∼** “intermediate sensitivity”

PDTOs with missing values were classified as “Unknown”.

### Bulk RNA sequencing of PDTOs

NucleoSpin RNA Plus (MN) kit was used to isolate total RNA from snap-frozen cell pellets. RNA samples were eluted in nuclease-free water and stored at -80 °C. Total RNA (100 ng) was ribo-depleted using Illumina’s Stranded Total RNA Prep with Ribo-Zero Plus (20040529) and library preparation was completed using standard Illumina protocols. Indexed libraries were pooled and sequenced on an Illumina NovaSeq6000 system (Illumina, San Diego, CA, USA), with one lane of the NovaSeq S4 flowcell (2x150 bp).

### Preprocessing PDTO bulk RNA-sequencing data

kallisto v.0.50.1 was used to extract transcript-level counts from fastq files. The human transcriptome index constructed from the Ensembl reference transcriptome version 108 was downloaded (https://github.com/pachterlab/kallisto-transcriptome-indices/releases) and the tximport R package v.1.32.0 was used to summarize transcript-level counts to gene-level and create a count matrix.

### Differential expression analysis (DEA) for PDTO bulk RNA sequencing data

The DESeq2 R package v1.40.2^14^ was used to perform DEA. Prefiltering was performed to exclude features with less than five normalized counts across n samples, where n equaled the lowest number of samples in the groups compared. Principal component analysis (PCA) revealed batch effects related to sequencing batch, but not anatomical site or passage number at time of RNA isolation. Therefore, sequencing batch was included as a covariate in the design formula. The biomaRt v.2.60.0 R package^15^ was used to convert Ensembl gene IDs to Entrez gene IDs and gene symbols. The EnhancedVolcano v.1.22.0 R package^16^ was used to create volcano plots to visualize DEA results.

### Pathway activity inference

The R implementation of the decoupleR framework (v.1.5.0)^17^ was used to infer pathway activities from bulk RNA sequencing and scRNA-seq data. The PROGENy^18^ weights of top 500 features were retrieved and the stat values from DEA results (DESeq2 output) were used as input for the multivariate linear model.

### Gene set enrichment analysis

For each DEA comparison, gene set enrichment analysis (GSEA) was performed with the fgsea^19^ method implemented in the clusterProfiler R package v.4.12.0^20^, using the curated (C2) and hallmark (H) gene sets available in the Human Molecular Signatures database (MSigDb)^21^. Pathways with Benjamini-Hochberg adjusted p values < 0.05 were considered significantly enriched.

### Molecular subtypes of PDTOs

The raw count matrix was filtered to exclude features with zero counts across all samples and variance stabilizing transformation was applied with DESeq2::vst() function with “blind=FALSE” setting. The IMF classification was implemented by using the CMScaller package^22^, using the transformed count matrix as input and the gene sets published by Joanito et al.^23^ as the template. The gene sets “iCMS2_Up” and “iCMS3_Down” were combined to form the iCMS2 class, while “iCMS2_Down” and “iCMS3_Up” were combined to form the iCMS3 class. PDTO samples were classified as iCMS2 or iCMS3 if the distance of a PDTO sample to the corresponding template was lower than that of 1,000 random permutations of the gene labels (FDR < 0.05). Samples with FDR values higher than the cutoff were defined as “Undefined”.

### Reanalysis of the CNP0004138 scRNA-seq dataset

Preprocessed scRNA-seq data of the CNP0004138 project was downloaded from the China National GeneBank Database^24^. The celltypist^25^ Python package (v.0.1.9) was used to annotate cell types by using the model “Immune_All_Low”. Cells annotated as “Epithelial cells” were retained for downstream analyses. For each treatment stage (pre-treatment and post-treatment), Seurat’s FindMarkers function was used to identify differentially expressed genes between clinical response groups. Differentially expressed genes overlapping with ISGs were visualized on a lollipop plot.

### RNA interference and drug treatment

HCT116 cells were grown in RPMI 1640 medium supplemented with 10% fetal bovine serum, 1% GlutaMAX, and 1% Penicillin-Streptomycin solution. Genomics Unit of Technology Center, Institute for Molecular Medicine Finland (FIMM) has authenticated the HCT116 cell line with the GenePrint24 system (Promega).

RNA interference was used to assess the role of *EPSTI1* in drug response. Briefly, 3 x 10^5^ HCT116 cells were plated in a 6-well plate, and on the next day, cells were transfected with Negative Control siRNA or EPSTI1 siRNA (#4392420, s41293, Ambion), using Lipofectamine RNAiMAX reagent (#13778030, Invitrogen). Seventy-two hours after transfection, cells were dissociated with TrypLE reagent (#12605028, Gibco), and 3 x 10^3^ cells were replated in a 96-well plate in four technical replicates. On the next day, cells were treated with fluorouracil or irinotecan, and 72 hours after drug treatment, alamarBlue cell viability reagent (A50100, Invitrogen) was used to measure cell viability.

### Statistical testing

Statistical analyses were performed on R version 4.4.1. The Wilcoxon rank-sum test was used to compare *in vitro* drug response estimates (AUC values) and gene module scores between two groups. The threshold for statistical significance was set to 0.05. Where appropriate, p values were adjusted with Benjamini-Hochberg or Bonferroni procedures.

## RESULTS

### *In vitro* modeling of drug responses with CRC PDTOs

Oncological management of patients with CRC typically involves a combination of conventional chemotherapeutics fluorouracil, irinotecan, and oxaliplatin. We hypothesized that integrating molecular profiles of primary CRC tumors and matched PDTOs with *in vitro* drug responses could reveal the molecular factors underlying heterogeneous responses to these drugs, thus allowing us to identify epithelial cancer cell-specific responses. To test this hypothesis, we used a subcohort of PDTOs established in the iCAN-PANORG project (Pan-cancer organoid biobank for precision medicine in abdominal cancers) and SYNCOPE (Systemic Neoadjuvant and adjuvant Control by Precision medicine in rectal cancer) trial (NCT04842006). PDTOs were established from tumor specimens of patients with colon or rectal adenocarcinoma (n = 75, **Methods**). Bulk RNA-seq was performed on treatment-naive PDTOs, and *in vitro* drug testing was performed in the next passage. Whole-exome sequencing data of the corresponding primary tumors was available for a subset of PDTOs (n = 44), which was used to infer the somatic mutations in PDTOs (**Figure 1A**). Data modalities and patient characteristics are presented in **Table 1**.

**Figure 1.**
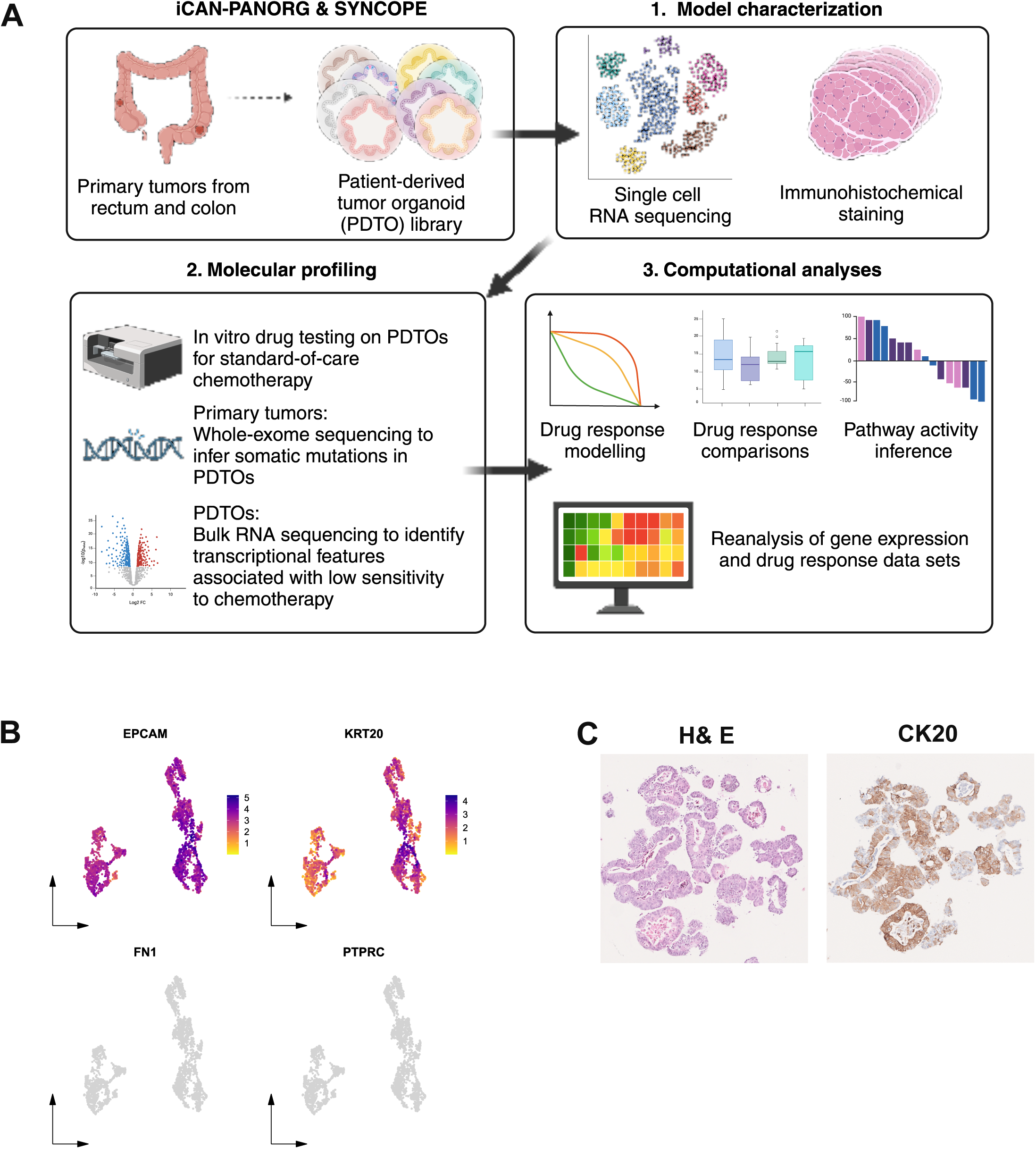
A. Overview of study design B. UMAP embedding of the scRNA-seq data of representative PDTOs. Genes marking epithelial cell identity (*EPCAM* and *KRT20*) are expressed at high levels in both PDTOs, whereas those marking immune cell (*PTPRC*) and stromal cells (*FN1*) have minimal expression. C. Immunohistochemical staining of representative PDTOs.

**Table 1.**
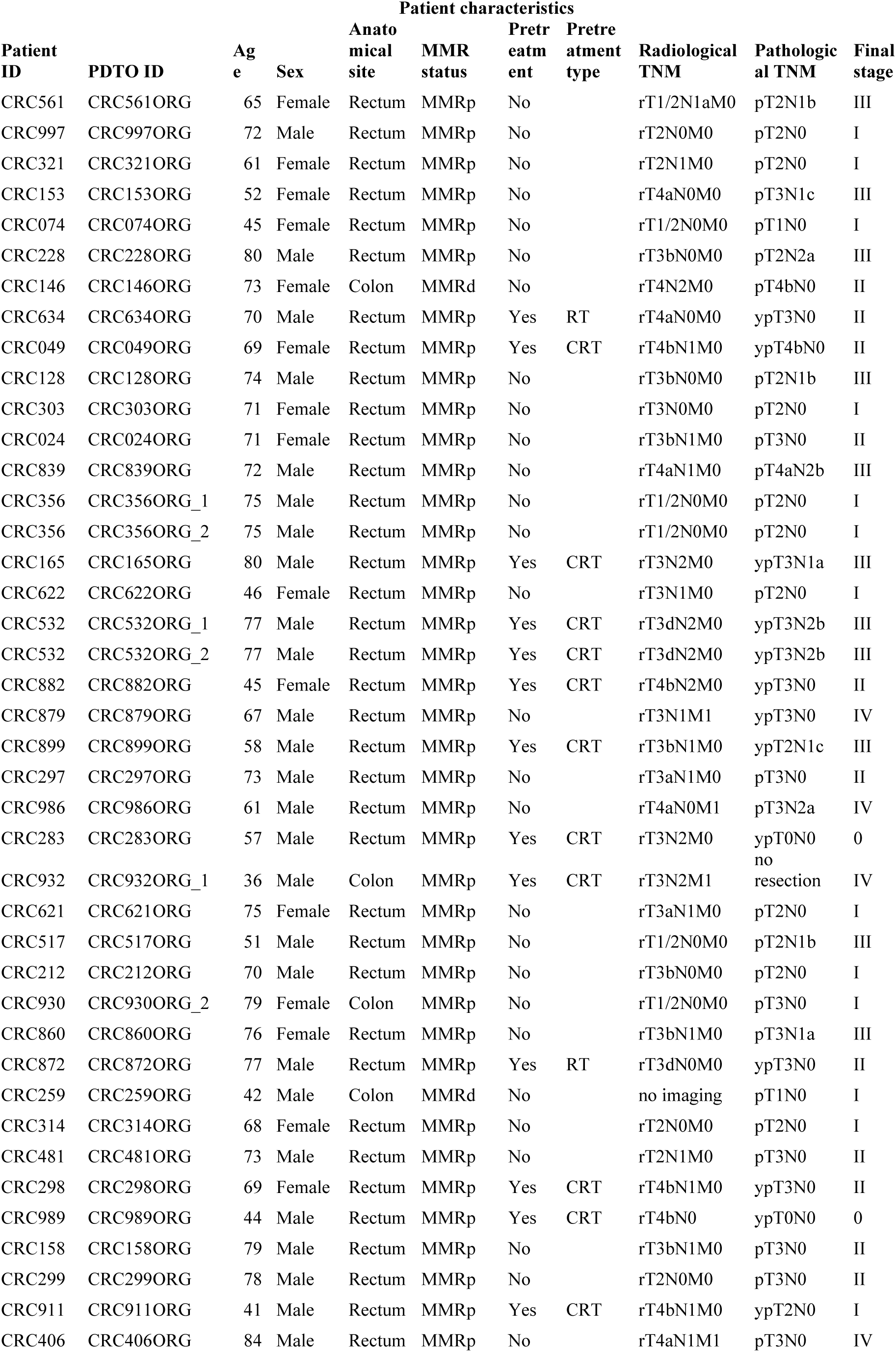
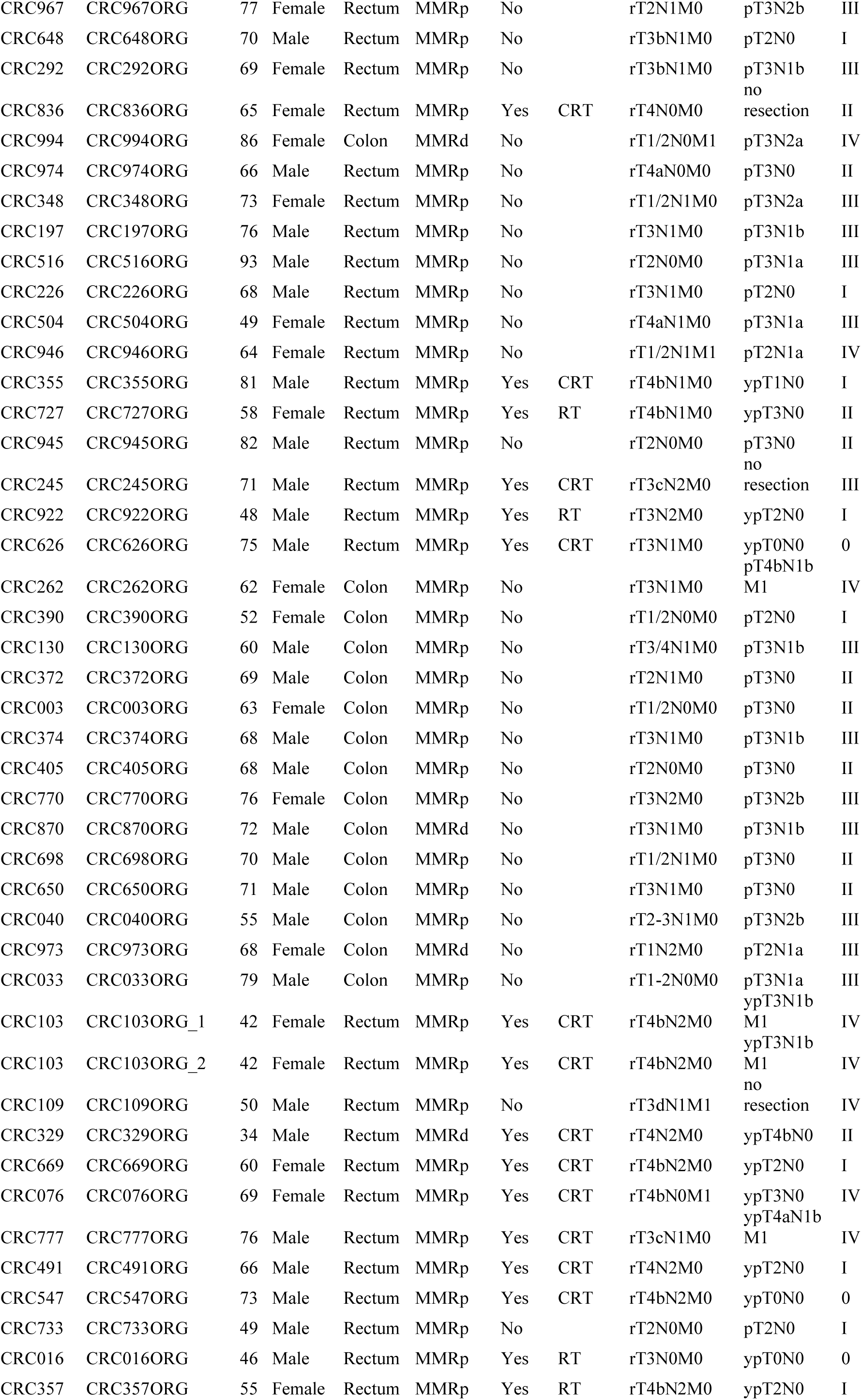
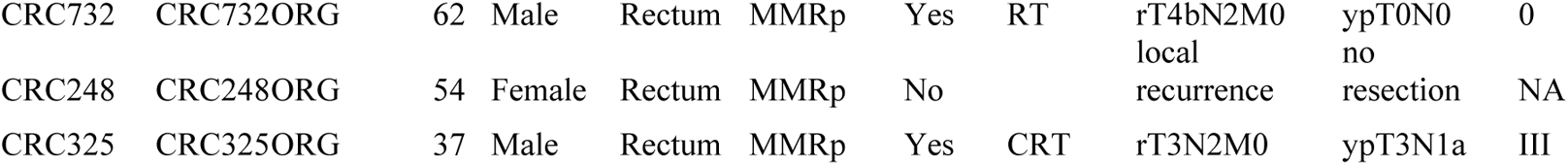
Overview of data modalities and patient characteristics. MMRp: Mismatch repair proficient, MMRd: Mismatch repair deficient, RT: Radiotherapy, CRT: Chemoradiotherapy.

The scRNA-seq data of two representative PDTOs revealed high *EPCAM* and *KRT20* expression and lack of gene expression marking immune (*PTPRC*) and stromal cells (*FN1*), confirming the epithelial nature of the PDTOs (**Figure 1B**). Immunohistochemical staining on representative PDTOs showed strong cytokeratin 20 expression, further confirming the epithelial nature of the models (**Figure 1C**).

We conducted *in vitro* drug testing on 75 PDTOs by treating them with fluorouracil, irinotecan, and oxaliplatin as single agents. PDTOs were treated with chemotherapeutics for five days, and dose-response curves were generated to calculate point estimates of drug responses (AUC values). After quality control (see **Methods**), 62 PDTOs had drug response data fulfilling the acceptance criteria. Drug testing on four representative PDTOs at two different time points showed high correlation coefficients between cell viability values, indicating the stability of *in vitro* drug responses across the time points tested (**Figure 2A**). *In vitro* drug responses of PDTOs to chemotherapeutics did not show strong correlations with each other (**Figure 2B**). Representative dose-response curves are shown in **Figure 2C**.

**Figure 2.**
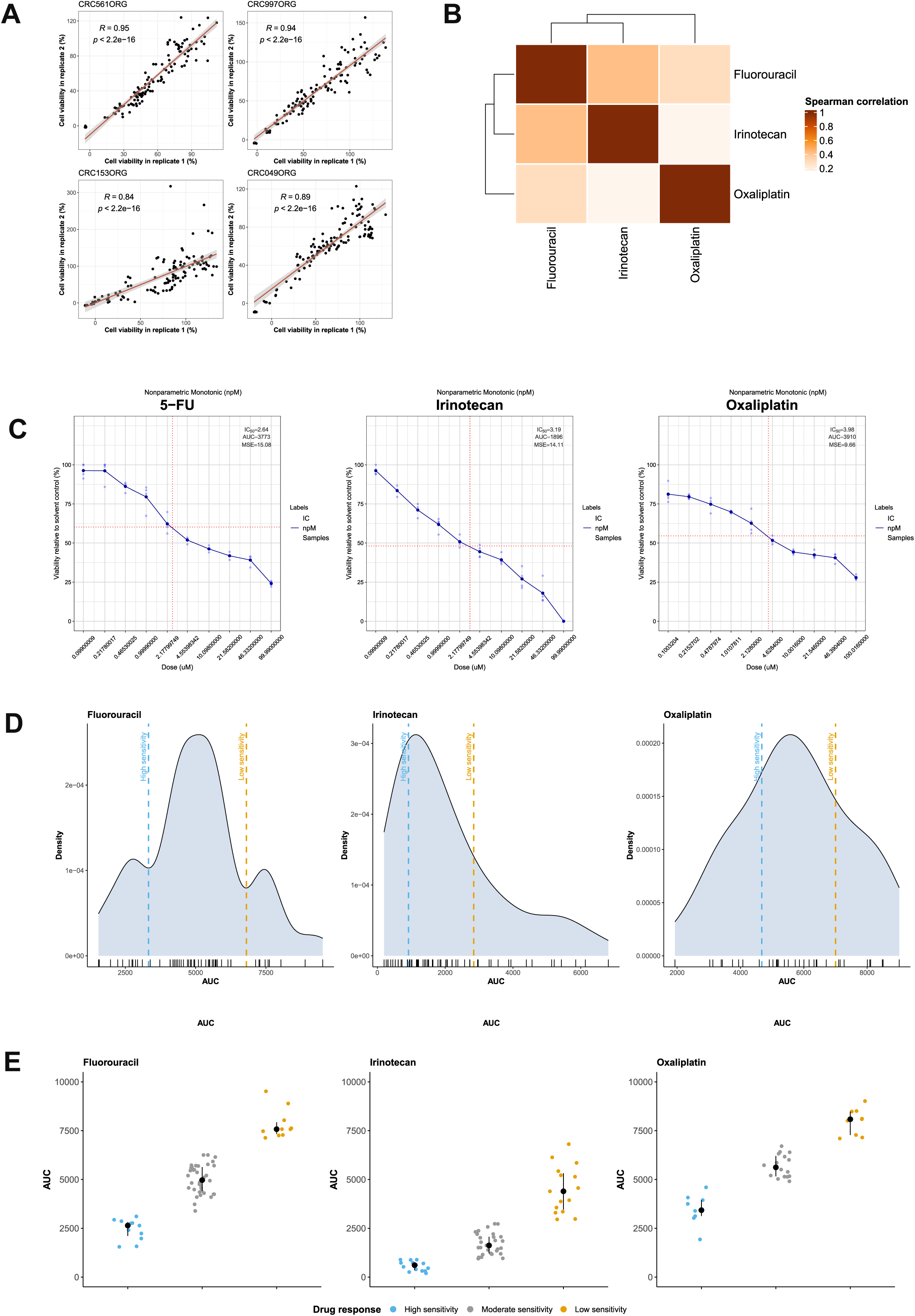
A. *In vitro* drug responses of PDTOs tested at different passages show highly correlated results. B. *In vitro* drug responses to standard-of-care chemotherapeutics do not show strong correlations. C. Representative dose-response curves for fluorouracil, irinotecan, and oxaliplatin. D. The distribution of AUC values are distinct for each chemotherapeutic agent. Dashed lines indicate classification thresholds. E. Dot plots showing AUC values with respect to *in vitro* drug response classification groups.

PDTOs displayed heterogeneous *in vitro* drug responses to chemotherapeutics, with irinotecan showing the highest efficacy (**Figure 2D**). Using the identified AUC thresholds (see **Methods**), we classified drug responses of PDTOs in low sensitivity, moderate sensitivity, and high sensitivity groups (**Figure 2E**), with the intention of achieving an analogous response classification mimicking no clinical response, partial response, and complete response categories.

### Somatic mutations and molecular subtypes associated with heterogeneous *in vitro* drug responses

We used the primary tumor WES data to infer the somatic mutations in PDTOs and compared *in vitro* drug responses between wild-type and mutant PDTOs. Of all genes tested, *CPEB2*, *CTNNB1*, *COL4A6*, *HLA-DRB5*, *IRS4*, and *TP53* mutations were associated with drug responses (BH-adjusted p-value < 0.05; **Supplementary Table 1**). Somatic *TP53* and *HLA-DRB5* mutations were associated with low sensitivity to fluorouracil, whereas somatic *CTNNB1* mutations were associated with higher sensitivity to both fluorouracil and oxaliplatin (**Figure 3A**).

**Figure 3.**
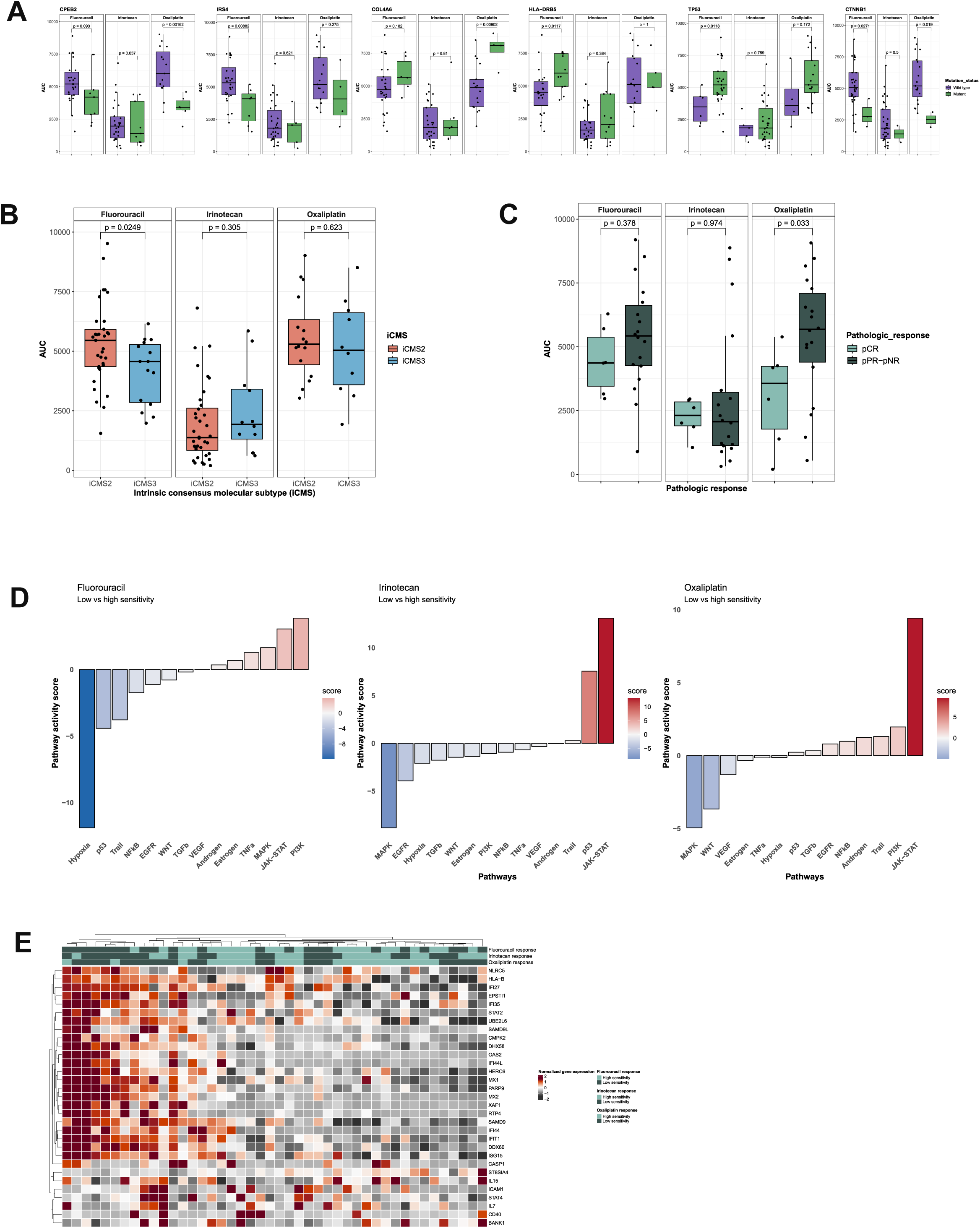
A. Box plots showing the impact of somatic mutations on *in vitro* drug responses to chemotherapeutics. AUC (*y* axis) values indicate in vitro drug responses, groups (*x* axis) indicate the inferred mutation status of PDTOs (wild type: purple, mutant: green). B. iCMS2 PDTOs show lower *in vitro* sensitivity to fluorouracil (*p* = 0.0249, Wilcoxon rank-sum test). C. Magnitude of *in vitro* response to chemotherapy stratified by tumor pathological response to neoadjuvant therapy. D. Waterfall plots showing differences in inferred pathway activities between PDTOs with low and high sensitivity to fluorouracil, irinotecan, and oxaliplatin. E. Heatmap showing ISG expression across CRC GDSC cell lines (n = 44). Notably, a subset of ISGs have higher expression in CRC cell lines with low sensitivity to chemotherapy.

Next, we asked whether molecular subtypes of PDTOs were associated with heterogeneous *in vitro* drug responses. The consensus molecular subtype (CMS) classification^26^ based on bulk-level gene expression profiles is not directly applicable to epithelial PDTOs, therefore we decided to use the epithelial cell-derived IMF classification system^23^. Using the CMScaller algorithm, we classified 36 PDTOs as iCMS2 and 15 PDTOs as iCMS3 (**Supplementary Table 2**). iCMS2 PDTOs had higher AUC for fluorouracil compared to iCMS3 PDTOs, indicating lower chemosensitivity (*p* = 0.0249, Wilcoxon rank-sum test) (**Figure 3B**). For irinotecan and oxaliplatin, the differences between the groups were not statistically significant (Irinotecan, *p* = 0.305; Oxaliplatin, *p* = 0.623).

To overcome the possible selection bias caused by studying only tumors with residual tumor after surgery, and to be able to characterize tumors that were potentially very responsive to therapy, we also compared *in vitro* drug responses of 32 PDTOs established from endoscopic rectal cancer biopsies taken before neoadjuvant therapy. Based on surgical pathologic response data, we classified PDTOs stratified to pathologic complete response (pCR) and incomplete response (pPR-pNR) of the primary tumors. For oxaliplatin, PDTOs derived from endoscopic biopsies of patients with pCR had lower AUC values compared to those with pPR-pNR, suggesting that PDTO- based drug testing can be used to model the efficacy of oxaliplatin *in vitro* (**Figure 3C**). In the case of irinotecan, the AUC values showed marginal differences between the groups, which can be attributed to the high efficacy of the compound *in vitro*.

### Epithelial Interferon signaling is associated with lower *in vitro* sensitivity to chemotherapy

To identify transcriptomic features associated with poor *in vitro* drug responses, we performed differential expression analysis (DEA) between 1) PDTOs with low and high chemosensitivity and 2) PDTOs with moderate and high chemosensitivity (**Methods**) (**Supplementary Figure 1A**). Then, we used multivariate linear modeling to infer pathway activities based on differentially expressed genes between the groups.

JAK-STAT pathway was ranked as the most active pathway in PDTOs with low chemosensitivity to irinotecan and oxaliplatin, and the second most active pathway in PDTOs with low chemosensitivity to fluorouracil (**Figure 3D**), possibly resulting from high expression of interferon-stimulated genes (ISGs) including *IFI6*, *IFIT1*, *IFIT2*, *IFIT3*, *ISG15*, and *MX1* (**Supplementary Figure 1B**). In parallel, we performed gene set enrichment analysis (GSEA) for hallmark gene sets (H) and curated gene sets (C2) in the MSigDB to identify transcriptional programs associated with low sensitivity to each chemotherapeutic. Consistent with high JAK-STAT pathway activity, hallmark gene sets for interferon alpha and interferon gamma response were ranked among the top gene sets in PDTOs with low sensitivity to irinotecan and oxaliplatin (**Supplementary Table 3**).

The lack of public databases linking gene expression profiles of PDTOs to their drug sensitivity profiles prompted us to look at the Genomics of Drug Sensitivity in Cancer (GDSC) database^27,28^, which provides gene expression and drug sensitivity data of cell line models of CRC. Focusing on 44 CRC cell lines with available gene expression and drug sensitivity data, we used the median IC50 values as a threshold to classify the cell lines having low or high sensitivity to fluorouracil, irinotecan, and oxaliplatin. Multiple ISGs, including *STAT1*, *IFI6*, *IFIT1*, *IFIT2*, and *IFIT3*, had higher gene expression in cell lines with low sensitivity to chemotherapeutics (**Figure 3E**), further supporting the association between ISG expression and low sensitivity of epithelial cancer cells to chemotherapy.

### Sustained epithelial interferon signaling underlies partial clinical response to therapy

To analyze how interferon signaling is linked to chemotherapy and clinical response, we reanalyzed the CNP0004138 scRNA-seq data set originating from patient-matched rectal tumor tissue specimens collected before and after neoadjuvant chemotherapy without radiotherapy^24^. Except for *IFI27* and *HLA-B*, ISG expression was restricted to only a small percentage of epithelial cells (**Supplementary Figure 2**).

To assess the relationship between ISG expression and clinical response, we focused on post-treatment samples and performed DEA between 1) tumors with pNR and pCR, and 2) tumors with pPR and pCR. Compared to tumors with pCR, both pPR and pNR tumors had higher expression of ISGs, including *IFIT1*, *MX1*, *ISG15*, and *EPSTI1* (**Figures 4A-B**). Epithelial-stromal interaction 1 (*EPSTI1*) is an ISG implicated in EMT regulation^29^, cell migration and invasion, and anchorage-independent growth^30^. Before chemotherapy, EPSTI1^+^ cell abundance was similar across clinical response groups. Chemotherapy induced a marked reduction in EPSTI1^+^ cells exclusively in pCR tumors (**Figure 4C**). Comparison of pathway activities across clinical response groups revealed that tumor necrosis factor (TNF)-related apoptosis-inducing ligand (TRAIL) pathway was active in both EPSTI1^+^ and EPSTI1^-^ cells only in pCR tumors (**Figure 4D**), suggesting a potential mechanism for selective elimination of EPSTI1^+^ cells.

**Figure 4.**
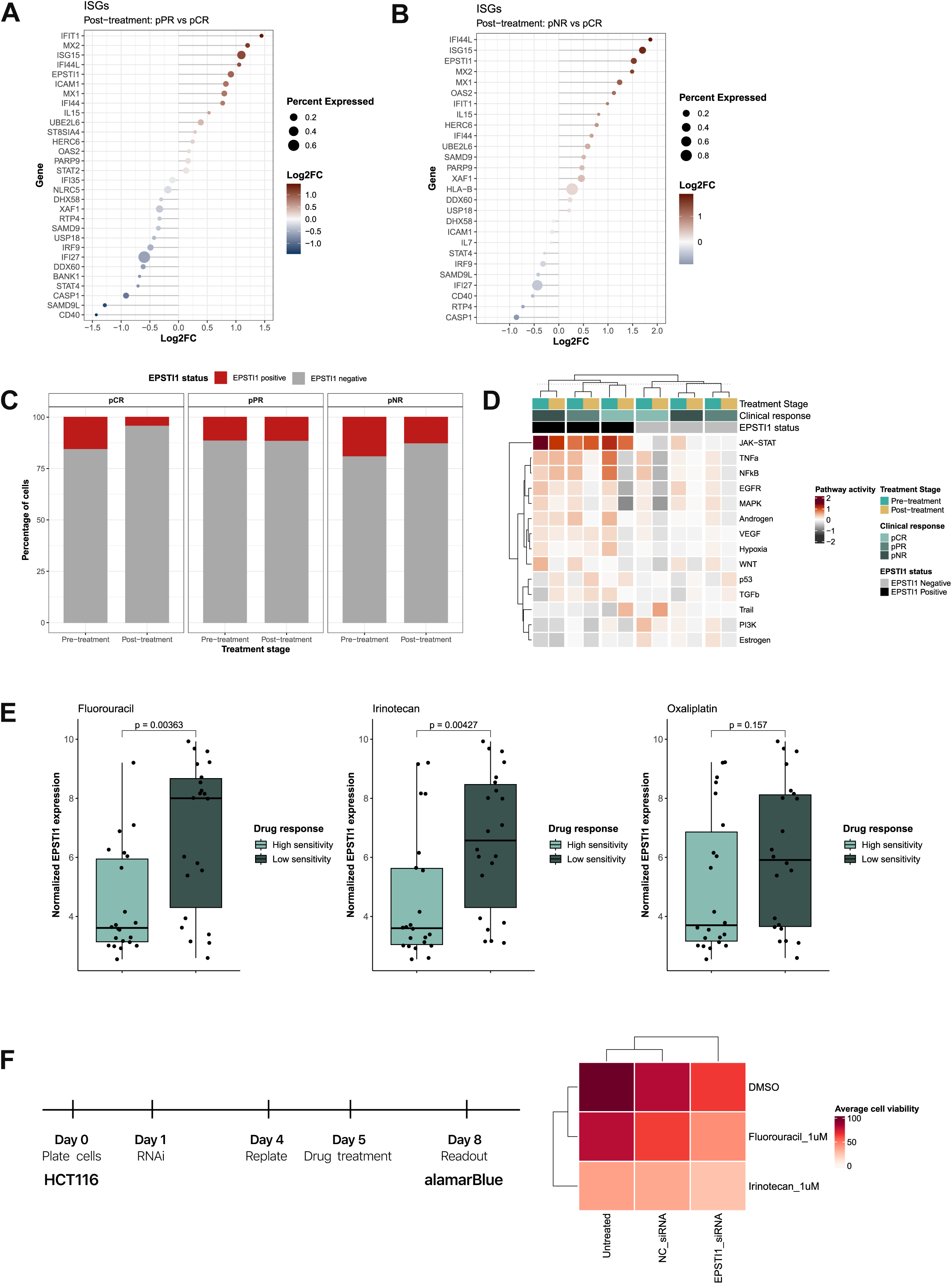
A-B. After chemotherapy, a subset of ISGs have higher expression in tumors with pPR and pNR. C. Chemotherapy leads to a sharp reduction in the fraction of EPSTI1^+^ cells in pCR tumors, but not in pPR and pNR tumors. D. Heatmap showing chemotherapy-induced changes in pathway activities with respect to EPSTI1 status and clinical response. pCR tumors have active Trail pathway, which distinguishes these tumors from pPR and pNR tumors. E. *EPSTI1* expression is higher in CRC cell lines with low *in vitro* sensitivity to chemotherapeutics (Fluorouracil, *p* = 0.00363; Irinotecan, *p* = 0.00427; Oxaliplatin, *p* = 0.00193, Wilcoxon rank-sum test). F. *EPSTI1* knockdown reduces cell viability and modulates response to chemotherapy.

To assess the link between *EPSTI1* and drug sensitivity, we used the GDSC data set and compared *EPSTI1* expression between CRC cells with high and low sensitivity to fluorouracil, irinotecan, and oxaliplatin. In all comparisons, *EPSTI1* expression was higher in CRC cell lines with low chemosensitivity (**Figure 4E**). RNA interference-mediated *EPSTI1* knockdown reduced CRC cell viability in the absence of chemotherapy and sensitized CRC cells to fluorouracil and irinotecan (**Figure 4F**). Collectively, these results suggest that sustained IFN signaling and IFN-responsive EPSTI1^+^ cells are likely to modulate incomplete pathologic response to chemotherapy.

## DISCUSSION

Building on our earlier work on pancreatic ductal adenocarcinoma PDTOs^31^, we aimed to utilize PDTOs as clinically relevant *in vitro* models of CRC to study epithelial cell-specific responses to chemotherapy. By combining genomic, transcriptomic, and phenotypic data of primary tumors and matched PDTOs, we identified genomic and transcriptomic factors underlying low *in vitro* drug sensitivity.

Integrating gene expression profiles of PDTOs and their *in vitro* drug responses, we identified several ISGs to be associated with lower sensitivity to chemotherapeutics, consistent with earlier reports linking IFN signaling to cancer stemness and drug resistance^32,33^. GDSC data set combining gene expression and drug response data of CRC cell lines provided supporting evidence to these observations. Single cell analyses of CRC tumors revealed chemotherapy-induced expression of ISGs, including *EPSTI1*, in tumors with incomplete pathologic response. While these observations link sustained interferon signaling and EPSTI1 to incomplete pathologic responses, it remains unclear whether EPSTI1 actively drives chemoresistance or is a byproduct of sustained IFN signaling.

Studies on *in vitro* models of CRC revealed translational regulation of *EPSTI1* expression by the KSR1/Erk signaling axis, which associates with the metastatic potential of cancer cells through regulation of cell mobility and EMT-like phenotype^30^. Pharmacological inhibition of ERK signaling suppresses EPSTI1 protein expression without altering transcript levels and can be used as a complementary strategy to target EPSTI1^+^ cells. Given chemotherapy-induced *EPSTI1* expression and the retention of EPSTI1^+^ cells in tumors with pPR and pNR, we hypothesize that this cell population plays a role in chemoresistance. While our preliminary results point out to EPSTI1’s role in cell survival and modulation of chemosensitivity, further studies on PDTO models of CRC will be necessary to portray the precise contribution of EPSTI1^+^ cells to CRC biology and clinical outcomes.

Earlier studies exploring genotype-phenotype relationships in PDTO models of CRC have linked *KRAS* mutations to the lack of response to the EGFR inhibitor cetuximab^34,35^. Our work, in contrast, aimed to identify putative associations between somatic mutations and *in vitro* drug responses to standard-of-care chemotherapy. Our observations linking somatic *TP53* and *CTNNB1* mutations to differential *in vitro* drug sensitivities support the reported information in the Catalogue of Somatic Mutations in Cancer (COSMIC) database. For other genes where no drug sensitivity/resistance data exists, gain- or loss-of-function studies on biologically and clinically relevant *in vitro* models of CRC (e.g., PDTOs) will be necessary to confirm the observed genotype-phenotype associations.

We acknowledge that the lack of tumor microenvironment (TME) is a well-recognized limitation of PDTOs, and *in vitro* drug responses of epithelial tumor cells cannot fully recapitulate the complex clinical response in vivo. However, PDTOs represent a biologically and clinically representative *in vitro* tumor model equipped to help identify how epithelial cancer cells respond to chemotherapeutics, which was this study’s main goal. The large-scale tissue acquisition for characterizing primary tumors and establishing PDTOs mainly from surgical specimens may cause selection bias to only focus on tumors without complete clinical and pathologic response and causes a lack of comparison to most responsive tumor subtypes both *in vivo* and *in vitro*. Another limitation is culture-associated transcriptional remodeling, a phenomenon that occurs over long-term culture^36^, which may confound the DEA and GSEA results. Temporal gene expression profiling (before and after drug treatment) on a larger set of PDTOs may help pinpoint dynamic pathway activations underlying drug responses and identify molecular vulnerabilities to be used for therapeutic intervention.

## CONCLUSION

Our study utilizing PDTO models offers new insights that contribute to explaining how epithelial cancer cells in CRC respond to chemotherapy, emphasizing the crucial role of JAK-STAT pathway and epithelial IFN signaling in incomplete pathologic response. Future studies aiming pharmaceutical inhibition of JAK-STAT or IFN signaling in preclinical CRC models are necessary to evaluate the therapeutic potential of these strategies.

## ACKNOWLEDGMENTS

The iCAN Flagship Project is funded by the Research Council of Finland (grant number 346555), University of Helsinki, Helsinki University Hospital, and consortium partner Boehringer Ingelheim. High-throughput sequencing of the primary tumors was performed at the Institute for Molecular Medicine Finland FIMM Genomics Unit and Biomedicum Functional Genomics Unit supported by HiLIFE and Biocenter Finland. Primary high-throughput sequencing data analyses were performed by FIMM Bioinformatics Unit supported by HiLIFE and Biocenter Finland.

T.T.S. was supported by funding from the Research Council of Finland and iCAN Digital Precision Cancer Medicine Flagship, and research grants by Jane and Aatos Erkko Foundation, Sigrid Juselius Foundation, Mary and Georg Ehrnrooth Foundation, Cancer Foundation Finland, Relander Foundation, and HUS and Pirha state research funding. The funding bodies had no role in drafting the article. RNA sequencing of PDTOs was performed at the Biomedicum Functional Genomics Unit at the Helsinki Institute of Life Science and Biocenter Finland at the University of Helsinki. Sequencing of scRNA-seq libraries was performed at Novogene Laboratories.

ICAN Flagship group authors are presented in the supplementary document.

## Abbreviations

AUC: Area under the curve
pCR: Pathologic complete response
CRC: Colorectal cancer
COSMIC: Catalogue of Somatic Mutations in Cancer
IFN: Interferon
iCMS: Intrinsic consensus molecular subtype
GDSC: Genomics of Drug Sensitivity in Cancer
GSEA: Gene set enrichment analysis
KDE: Kernel density estimation
MSigDB: Molecular Signatures Database
pNR: Pathologic no response
pPR: Pathologic partial response
PDTO: Patient-derived tumor organoid
scRNA-seq: Single-cell RNA sequencing
TME: Tumor microenvironment
UMAP: Uniform manifold approximation and projection

## Author contributions

**Erdogan Pekcan Erkan:** Investigation, Formal analysis, Validation, Data Curation, Writing – Original Draft, Visualization, Writing – Review & Editing **Emmi Hämäläinen**: Software, Data Curation, Formal analysis, Visualization, Writing – Review & Editing **Julia Kolikova:** Investigation **Kornelia Kuc:** Investigation **Kalle Ojala**: Data curation **Meiju Kukkonen:** Investigation **Ismail Hermelo:** Investigation **Selja Koskensalo:** Resources **Alli Leppä:** Resources **Ilona Keränen:** Resources **Carola Haapamäki:** Resources **Essi Karjalainen:** Resources **Monica Carpelan- Holmström:** Resources **Laura Renkonen-Sinisalo:** Resources **Laura Koskenvuo:** Resources **Pauli Puolakkainen:** Resources **Jukka-Pekka Mecklin:** Resources, Writing – Review & Editing **Elina Virolainen:** Resources **Anna Lepistö:** Resources, Writing – Review & Editing **iCAN:** Resources **Matti Nykter:** Supervision, Writing – Review & Editing **Lauri A. Aaltonen:** Resources **Ari Ristimäk**i: Resources, Writing – Review & Editing **Toni T. Seppälä:** Conception and design, Supervision, Data curation, Resources, Project administration, Funding acquisition, Writing – Review & Editing

## Data transparency statement

Processed bulk RNA-seq will be deposited in Gene Expression Omnibus and made publicly available at the time of publication. All computational codes will be deposited to Zenodo and made publicly available at the time of publication. Additional information required to reanalyze the data will be provided by the lead contact.

---

**Supplementary Figure 1.**
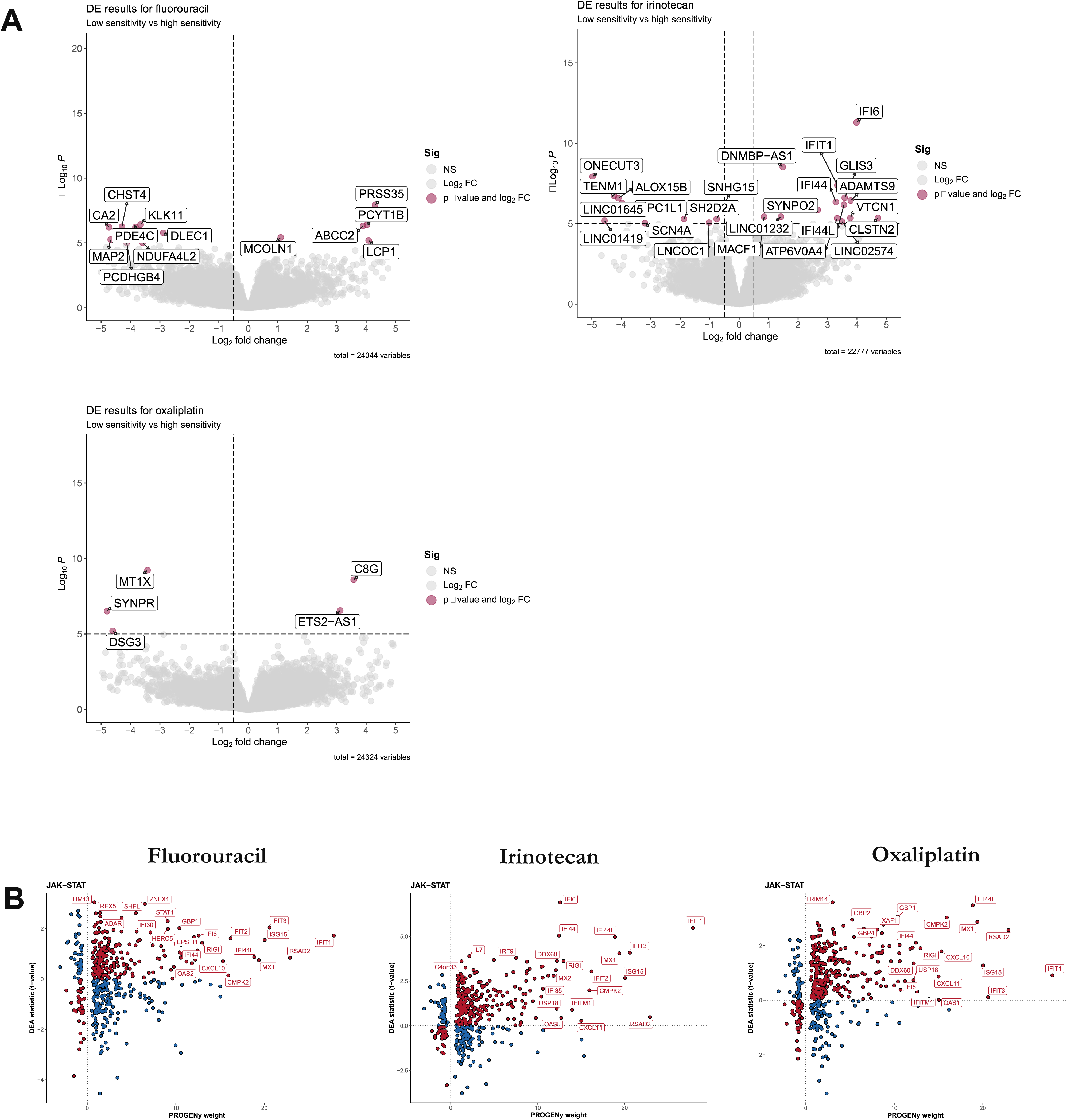
A. Volcano plots showing DEA results. B. Scatter plots showing the relationship between JAK-STAT target gene weights and corresponding t-values calculated in DEA for each comparison.

**Supplementary Figure 2.**
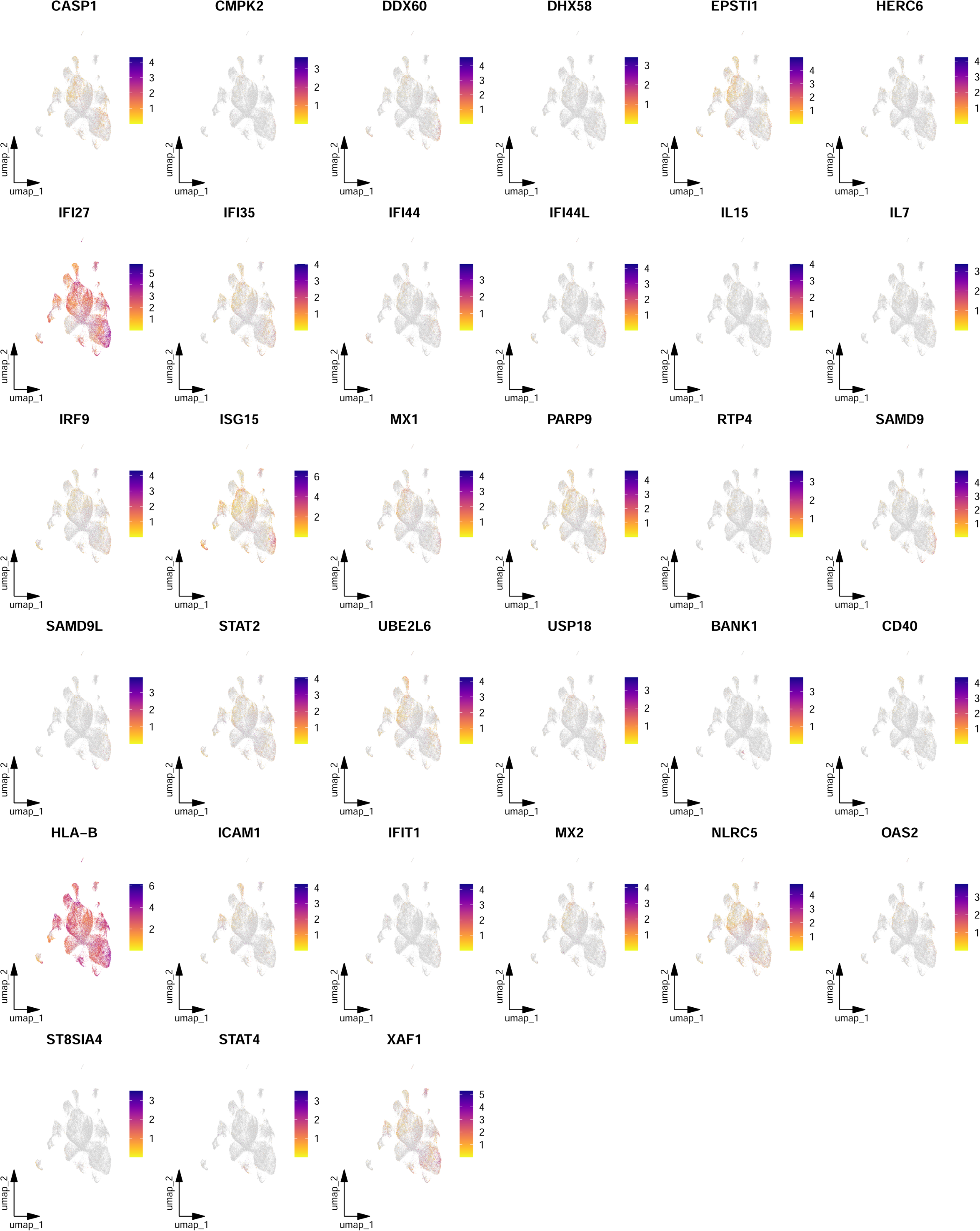
Feature plots showing ISG expression in epithelial cells. Colors indicate gene expression.

**Supplementary Table 1.** Somatic mutations associated with differential in vitro drug responses.

| gene_symbol | drug | not_mutated_n | mutated_n | p_value | adjusted_p_value | is_significant |
| --- | --- | --- | --- | --- | --- | --- |
| CPEB2 | Oxaliplatin | 6 | 15 | 0,0016 | 0,00486 | yes |
| IRS4 | Fluorouracil | 7 | 28 | 0,0088 | 0,02646 | yes |
| COL4A6 | Oxaliplatin | 4 | 17 | 0,009 | 0,02706 | yes |
| HLA-DRB5 | Fluorouracil | 12 | 23 | 0,0117 | 0,0351 | yes |
| TP53 | Fluorouracil | 29 | 6 | 0,0118 | 0,0354 | yes |
| CTNNB1 | Fluorouracil | 3 | 32 | 0,0271 | 0,04065 | yes |
| CTNNB1 | Oxaliplatin | 2 | 19 | 0,019 | 0,04065 | yes |
| SAFB2 | Fluorouracil | 7 | 28 | 0,0177 | 0,0531 | no |
| MYO16 | Fluorouracil | 10 | 25 | 0,0185 | 0,0555 | no |
| RP1L1 | Fluorouracil | 6 | 29 | 0,0242 | 0,0726 | no |
| YBX1P6 | Irinotecan | 7 | 28 | 0,0304 | 0,0912 | no |
| H1-8 | Irinotecan | 5 | 30 | 0,0403 | 0,1209 | no |
| DTD2 | Fluorouracil | 7 | 28 | 0,0665 | 0,1305 | no |
| DTD2 | Irinotecan | 7 | 28 | 0,087 | 0,1305 | no |
| MUC6 | Oxaliplatin | 13 | 8 | 0,0446 | 0,1338 | no |
| CPEB2 | Fluorouracil | 9 | 26 | 0,093 | 0,1395 | no |
| SDK1 | Fluorouracil | 6 | 29 | 0,0693 | 0,165 | no |
| SDK1 | Irinotecan | 6 | 29 | 0,11 | 0,165 | no |
| RNF213 | Irinotecan | 8 | 27 | 0,0567 | 0,168 | no |
| RNF213 | Oxaliplatin | 6 | 15 | 0,112 | 0,168 | no |
| ANK2 | Irinotecan | 7 | 28 | 0,122 | 0,183 | no |
| ANK2 | Oxaliplatin | 4 | 17 | 0,0805 | 0,183 | no |
| RYR1 | Irinotecan | 13 | 22 | 0,0678 | 0,2034 | no |
| LMTK3 | Fluorouracil | 5 | 30 | 0,0695 | 0,2085 | no |
| ERBB2 | Irinotecan | 4 | 31 | 0,139 | 0,2085 | no |
| ERBB2 | Oxaliplatin | 3 | 18 | 0,101 | 0,2085 | no |
| IGLV5-45 | Irinotecan | 7 | 28 | 0,0728 | 0,2184 | no |
| AR | Fluorouracil | 5 | 30 | 0,0771 | 0,2313 | no |
| TGFBR2 | Fluorouracil | 5 | 30 | 0,0771 | 0,2313 | no |
| ACVR2A | Fluorouracil | 5 | 30 | 0,0771 | 0,2313 | no |
| FBXW7 | Fluorouracil | 9 | 26 | 0,0792 | 0,2376 | no |
| MT-ND5 | Fluorouracil | 6 | 29 | 0,0838 | 0,2514 | no |
| SACS | Fluorouracil | 6 | 29 | 0,0838 | 0,2514 | no |
| FBXO11 | Fluorouracil | 7 | 28 | 0,257 | 0,257 | no |
| FBXO11 | Irinotecan | 6 | 29 | 0,246 | 0,257 | no |
| FBXO11 | Oxaliplatin | 4 | 17 | 0,12 | 0,257 | no |
| TP53 | Oxaliplatin | 17 | 4 | 0,172 | 0,258 | no |
| PIK3CA | Fluorouracil | 7 | 28 | 0,087 | 0,261 | no |
| COL4A6 | Fluorouracil | 6 | 29 | 0,182 | 0,273 | no |
| HTT | Irinotecan | 9 | 26 | 0,093 | 0,279 | no |
| HRNR | Fluorouracil | 9 | 26 | 0,146 | 0,2895 | no |
| HRNR | Irinotecan | 9 | 26 | 0,193 | 0,2895 | no |
| ZNF831 | Fluorouracil | 5 | 30 | 0,104 | 0,312 | no |
| CHEK2 | Fluorouracil | 6 | 29 | 0,212 | 0,318 | no |
| CHEK2 | Irinotecan | 5 | 30 | 0,138 | 0,318 | no |
| KRAS | Fluorouracil | 8 | 27 | 0,216 | 0,324 | no |
| KRAS | Oxaliplatin | 6 | 15 | 0,132 | 0,324 | no |
| CPAMD8 | Oxaliplatin | 5 | 16 | 0,109 | 0,327 | no |
| PCDH10 | Irinotecan | 4 | 31 | 0,223 | 0,3345 | no |
| PCDH10 | Oxaliplatin | 2 | 19 | 0,114 | 0,3345 | no |
| ANKRD36C | Fluorouracil | 7 | 28 | 0,112 | 0,336 | no |
| CPAMD8 | Fluorouracil | 6 | 29 | 0,229 | 0,3435 | no |
| RYR2 | Irinotecan | 6 | 29 | 0,168 | 0,3555 | no |
| RYR2 | Oxaliplatin | 4 | 17 | 0,237 | 0,3555 | no |
| CFAP46 | Fluorouracil | 6 | 29 | 0,12 | 0,36 | no |
| HNF1A | Irinotecan | 6 | 29 | 0,12 | 0,36 | no |
| SHANK3 | Irinotecan | 7 | 28 | 0,122 | 0,366 | no |
| IGLV5-45 | Fluorouracil | 8 | 27 | 0,246 | 0,369 | no |
| LRP2 | Fluorouracil | 5 | 30 | 0,125 | 0,375 | no |
| KIF26A | Irinotecan | 5 | 30 | 0,125 | 0,375 | no |
| UBR5 | Fluorouracil | 5 | 30 | 0,125 | 0,375 | no |
| CACNA1E | Fluorouracil | 5 | 30 | 0,125 | 0,375 | no |
| GDF11 | Oxaliplatin | 3 | 18 | 0,125 | 0,375 | no |
| FAT4 | Irinotecan | 12 | 23 | 0,131 | 0,393 | no |
| TIAM1 | Fluorouracil | 6 | 29 | 0,264 | 0,396 | no |
| TIAM1 | Irinotecan | 6 | 29 | 0,155 | 0,396 | no |
| SHANK3 | Fluorouracil | 7 | 28 | 0,353 | 0,398 | no |
| SHANK3 | Oxaliplatin | 5 | 16 | 0,398 | 0,398 | no |
| SDK1 | Oxaliplatin | 5 | 16 | 0,398 | 0,398 | no |
| ZFHX4 | Fluorouracil | 3 | 32 | 0,133 | 0,399 | no |
| MYO16 | Irinotecan | 9 | 26 | 0,266 | 0,399 | no |
| FAT4 | Fluorouracil | 13 | 22 | 0,282 | 0,4 | no |
| FAT4 | Oxaliplatin | 7 | 14 | 0,4 | 0,4 | no |
| NBEA | Fluorouracil | 10 | 25 | 0,221 | 0,4 | no |
| NBEA | Irinotecan | 10 | 25 | 0,371 | 0,4 | no |
| NBEA | Oxaliplatin | 7 | 14 | 0,4 | 0,4 | no |
| ASCL1 | Fluorouracil | 6 | 29 | 0,347 | 0,412 | no |
| ASCL1 | Irinotecan | 5 | 30 | 0,248 | 0,412 | no |
| ASCL1 | Oxaliplatin | 3 | 18 | 0,412 | 0,412 | no |
| IRS4 | Oxaliplatin | 4 | 17 | 0,275 | 0,4125 | no |
| RP1L1 | Oxaliplatin | 4 | 17 | 0,275 | 0,4125 | no |
| PAK2 | Fluorouracil | 11 | 24 | 0,14 | 0,42 | no |
| POTEI | Fluorouracil | 6 | 29 | 0,284 | 0,426 | no |
| POTEI | Oxaliplatin | 4 | 17 | 0,172 | 0,426 | no |
| ABCF1 | Fluorouracil | 14 | 21 | 0,143 | 0,429 | no |
| ATN1 | Irinotecan | 7 | 28 | 0,143 | 0,429 | no |
| PCDH10 | Fluorouracil | 5 | 30 | 0,437 | 0,437 | no |
| RIMBP2 | Fluorouracil | 9 | 26 | 0,146 | 0,438 | no |
| DSCAML1 | Fluorouracil | 7 | 28 | 0,208 | 0,445 | no |
| DSCAML1 | Irinotecan | 7 | 28 | 0,312 | 0,445 | no |
| DSCAML1 | Oxaliplatin | 5 | 16 | 0,445 | 0,445 | no |
| BLTP1 | Fluorouracil | 6 | 29 | 0,155 | 0,465 | no |
| FBXW7 | Irinotecan | 8 | 27 | 0,467 | 0,467 | no |
| FBXW7 | Oxaliplatin | 8 | 13 | 0,456 | 0,467 | no |
| MUC4 | Fluorouracil | 16 | 19 | 0,159 | 0,477 | no |
| ABCA13 | Fluorouracil | 10 | 25 | 0,315 | 0,483 | no |
| ABCA13 | Oxaliplatin | 7 | 14 | 0,322 | 0,483 | no |
| RBMXL1 | Irinotecan | 6 | 29 | 0,325 | 0,4875 | no |
| RBMXL1 | Oxaliplatin | 4 | 17 | 0,275 | 0,4875 | no |
| FRMPD4 | Irinotecan | 5 | 30 | 0,164 | 0,492 | no |
| YBX1P6 | Fluorouracil | 7 | 28 | 0,496 | 0,496 | no |
| YBX1P6 | Oxaliplatin | 3 | 18 | 0,412 | 0,496 | no |
| CTNNB1 | Irinotecan | 2 | 33 | 0,5 | 0,5 | no |
| LCORL | Irinotecan | 7 | 28 | 0,167 | 0,501 | no |
| CSMD3 | Fluorouracil | 7 | 28 | 0,274 | 0,5145 | no |
| CSMD3 | Oxaliplatin | 2 | 19 | 0,343 | 0,5145 | no |
| TIAM1 | Oxaliplatin | 4 | 17 | 0,517 | 0,517 | no |
| ABCF1 | Oxaliplatin | 9 | 12 | 0,345 | 0,5175 | no |
| CHEK2 | Oxaliplatin | 6 | 15 | 0,519 | 0,519 | no |
| RYR1 | Fluorouracil | 12 | 23 | 0,52 | 0,52 | no |
| RYR1 | Oxaliplatin | 6 | 15 | 0,424 | 0,52 | no |
| APC | Fluorouracil | 29 | 6 | 0,526 | 0,534 | no |
| APC | Irinotecan | 29 | 6 | 0,418 | 0,534 | no |
| APC | Oxaliplatin | 18 | 3 | 0,534 | 0,534 | no |
| MUC3A | Fluorouracil | 7 | 28 | 0,312 | 0,534 | no |
| MUC3A | Irinotecan | 7 | 28 | 0,496 | 0,534 | no |
| MUC3A | Oxaliplatin | 3 | 18 | 0,534 | 0,534 | no |
| ZC3H13 | Fluorouracil | 6 | 29 | 0,47 | 0,534 | no |
| ZC3H13 | Irinotecan | 5 | 30 | 0,465 | 0,534 | no |
| ZC3H13 | Oxaliplatin | 3 | 18 | 0,534 | 0,534 | no |
| CSMD1 | Fluorouracil | 7 | 28 | 0,18 | 0,54 | no |
| ANK3 | Fluorouracil | 3 | 32 | 0,361 | 0,5415 | no |
| ANK3 | Oxaliplatin | 2 | 19 | 0,19 | 0,5415 | no |
| CFAP46 | Oxaliplatin | 4 | 17 | 0,362 | 0,543 | no |
| MT-ND1 | Fluorouracil | 4 | 31 | 0,551 | 0,551 | no |
| MT-ND1 | Irinotecan | 4 | 31 | 0,517 | 0,551 | no |
| MT-ND1 | Oxaliplatin | 3 | 18 | 0,534 | 0,551 | no |
| ANK3 | Irinotecan | 4 | 31 | 0,551 | 0,551 | no |
| MUC4 | Irinotecan | 15 | 20 | 0,377 | 0,554 | no |
| MUC4 | Oxaliplatin | 9 | 12 | 0,554 | 0,554 | no |
| ATM | Fluorouracil | 9 | 26 | 0,559 | 0,559 | no |
| ATM | Irinotecan | 9 | 26 | 0,462 | 0,559 | no |
| ATM | Oxaliplatin | 5 | 16 | 0,313 | 0,559 | no |
| DTD2 | Oxaliplatin | 4 | 17 | 0,574 | 0,574 | no |
| HLA-DRB5 | Irinotecan | 13 | 22 | 0,384 | 0,576 | no |
| ZFHX3 | Fluorouracil | 7 | 28 | 0,194 | 0,582 | no |
| MUC6 | Fluorouracil | 21 | 14 | 0,391 | 0,5865 | no |
| HTT | Fluorouracil | 9 | 26 | 0,396 | 0,594 | no |
| RUNX1 | Fluorouracil | 5 | 30 | 0,556 | 0,6 | no |
| RUNX1 | Irinotecan | 5 | 30 | 0,524 | 0,6 | no |
| RUNX1 | Oxaliplatin | 3 | 18 | 0,6 | 0,6 | no |
| H1-8 | Oxaliplatin | 2 | 19 | 0,4 | 0,6 | no |
| KRTAP13-3 | Fluorouracil | 16 | 19 | 0,227 | 0,602 | no |
| KRTAP13-3 | Irinotecan | 16 | 19 | 0,518 | 0,602 | no |
| KRTAP13-3 | Oxaliplatin | 12 | 9 | 0,602 | 0,602 | no |
| ANK2 | Fluorouracil | 7 | 28 | 0,606 | 0,606 | no |
| ANKRD36C | Irinotecan | 7 | 28 | 0,606 | 0,606 | no |
| ANKRD36C | Oxaliplatin | 4 | 17 | 0,41 | 0,606 | no |
| DNAH5 | Fluorouracil | 7 | 28 | 0,606 | 0,606 | no |
| DNAH5 | Irinotecan | 8 | 27 | 0,231 | 0,606 | no |
| DNAH5 | Oxaliplatin | 4 | 17 | 0,574 | 0,606 | no |
| SELENOP | Fluorouracil | 16 | 19 | 0,608 | 0,608 | no |
| SELENOP | Irinotecan | 15 | 20 | 0,537 | 0,608 | no |
| SELENOP | Oxaliplatin | 9 | 12 | 0,602 | 0,608 | no |
| CACNA1E | Irinotecan | 5 | 30 | 0,409 | 0,6135 | no |
| ZNF831 | Oxaliplatin | 4 | 17 | 0,41 | 0,615 | no |
| LMTK3 | Oxaliplatin | 3 | 18 | 0,412 | 0,618 | no |
| IRS4 | Irinotecan | 5 | 30 | 0,621 | 0,621 | no |
| HSPD1 | Fluorouracil | 11 | 24 | 0,207 | 0,621 | no |
| DNAH10 | Fluorouracil | 4 | 31 | 0,392 | 0,622 | no |
| DNAH10 | Irinotecan | 4 | 31 | 0,622 | 0,622 | no |
| DNAH10 | Oxaliplatin | 2 | 19 | 0,533 | 0,622 | no |
| PRRC2A | Oxaliplatin | 5 | 16 | 0,208 | 0,624 | no |
| TCF7L2 | Fluorouracil | 8 | 27 | 0,421 | 0,6315 | no |
| TCF7L2 | Irinotecan | 10 | 25 | 0,391 | 0,6315 | no |
| CPEB2 | Irinotecan | 9 | 26 | 0,637 | 0,637 | no |
| CPAMD8 | Irinotecan | 6 | 29 | 0,646 | 0,646 | no |
| POTEI | Irinotecan | 5 | 30 | 0,654 | 0,654 | no |
| NRAS | Fluorouracil | 5 | 30 | 0,437 | 0,6555 | no |
| NRAS | Irinotecan | 6 | 29 | 0,304 | 0,6555 | no |
| KL | Fluorouracil | 11 | 24 | 0,582 | 0,657 | no |
| KL | Irinotecan | 11 | 24 | 0,657 | 0,657 | no |
| KL | Oxaliplatin | 9 | 12 | 0,554 | 0,657 | no |
| LMTK3 | Irinotecan | 4 | 31 | 0,659 | 0,659 | no |
| APOB | Fluorouracil | 6 | 29 | 0,444 | 0,666 | no |
| APOB | Oxaliplatin | 4 | 17 | 0,317 | 0,666 | no |
| ZNF469 | Fluorouracil | 4 | 31 | 0,223 | 0,669 | no |
| MSH6 | Fluorouracil | 4 | 31 | 0,223 | 0,669 | no |
| MUC6 | Irinotecan | 20 | 15 | 0,677 | 0,677 | no |
| KCNT1 | Fluorouracil | 7 | 28 | 0,496 | 0,678 | no |
| KCNT1 | Irinotecan | 6 | 29 | 0,678 | 0,678 | no |
| KCNT1 | Oxaliplatin | 4 | 17 | 0,317 | 0,678 | no |
| CCDC168 | Fluorouracil | 8 | 27 | 0,28 | 0,68 | no |
| CCDC168 | Irinotecan | 8 | 27 | 0,68 | 0,68 | no |
| CCDC168 | Oxaliplatin | 6 | 15 | 0,47 | 0,68 | no |
| CSMD3 | Irinotecan | 8 | 27 | 0,68 | 0,68 | no |
| MUC19 | Irinotecan | 16 | 19 | 0,227 | 0,681 | no |
| PKD1 | Fluorouracil | 4 | 31 | 0,392 | 0,686 | no |
| PKD1 | Irinotecan | 4 | 31 | 0,484 | 0,686 | no |
| PKD1 | Oxaliplatin | 2 | 19 | 0,686 | 0,686 | no |
| RYR2 | Fluorouracil | 5 | 30 | 0,689 | 0,689 | no |
| KIF26A | Fluorouracil | 5 | 30 | 0,689 | 0,689 | no |
| KIF26A | Oxaliplatin | 5 | 16 | 0,495 | 0,689 | no |
| SP8 | Fluorouracil | 5 | 30 | 0,689 | 0,689 | no |
| SP8 | Irinotecan | 5 | 30 | 0,524 | 0,689 | no |
| SP8 | Oxaliplatin | 4 | 17 | 0,462 | 0,689 | no |
| LCORL | Fluorouracil | 7 | 28 | 0,695 | 0,695 | no |
| LCORL | Oxaliplatin | 4 | 17 | 0,574 | 0,695 | no |
| DCHS2 | Irinotecan | 5 | 30 | 0,465 | 0,6975 | no |
| DCHS2 | Oxaliplatin | 2 | 19 | 0,286 | 0,6975 | no |
| LTN1 | Fluorouracil | 5 | 30 | 0,311 | 0,6975 | no |
| LTN1 | Irinotecan | 5 | 30 | 0,465 | 0,6975 | no |
| FAT3 | Fluorouracil | 6 | 29 | 0,678 | 0,698 | no |
| FAT3 | Irinotecan | 6 | 29 | 0,646 | 0,698 | no |
| FAT3 | Oxaliplatin | 4 | 17 | 0,698 | 0,698 | no |
| MYO15B | Irinotecan | 4 | 31 | 0,392 | 0,7065 | no |
| MYO15B | Oxaliplatin | 3 | 18 | 0,471 | 0,7065 | no |
| DNAH14 | Fluorouracil | 4 | 31 | 0,364 | 0,7065 | no |
| DNAH14 | Oxaliplatin | 3 | 18 | 0,471 | 0,7065 | no |
| CACNA1E | Oxaliplatin | 5 | 16 | 0,719 | 0,719 | no |
| MT-ND5 | Irinotecan | 7 | 28 | 0,726 | 0,74 | no |
| MT-ND5 | Oxaliplatin | 3 | 18 | 0,74 | 0,74 | no |
| LTN1 | Oxaliplatin | 3 | 18 | 0,74 | 0,74 | no |
| NRAS | Oxaliplatin | 3 | 18 | 0,74 | 0,74 | no |
| ADAMTS20 | Irinotecan | 5 | 30 | 0,248 | 0,744 | no |
| CASP5 | Fluorouracil | 7 | 28 | 0,293 | 0,744 | no |
| CASP5 | Irinotecan | 7 | 28 | 0,496 | 0,744 | no |
| FREM2 | Fluorouracil | 6 | 29 | 0,497 | 0,7455 | no |
| FREM2 | Irinotecan | 6 | 29 | 0,418 | 0,7455 | no |
| PTEN | Fluorouracil | 3 | 32 | 0,498 | 0,747 | no |
| PTEN | Irinotecan | 3 | 32 | 0,361 | 0,747 | no |
| MYO16 | Oxaliplatin | 8 | 13 | 0,75 | 0,75 | no |
| TP53 | Irinotecan | 30 | 5 | 0,759 | 0,759 | no |
| FRG2C | Fluorouracil | 8 | 27 | 0,444 | 0,763 | no |
| FRG2C | Irinotecan | 8 | 27 | 0,623 | 0,763 | no |
| FRG2C | Oxaliplatin | 4 | 17 | 0,763 | 0,763 | no |
| ARID1A | Fluorouracil | 5 | 30 | 0,654 | 0,763 | no |
| ARID1A | Irinotecan | 5 | 30 | 0,437 | 0,763 | no |
| ARID1A | Oxaliplatin | 4 | 17 | 0,763 | 0,763 | no |
| CACNA1H | Fluorouracil | 6 | 29 | 0,743 | 0,763 | no |
| CACNA1H | Irinotecan | 5 | 30 | 0,759 | 0,763 | no |
| CACNA1H | Oxaliplatin | 4 | 17 | 0,763 | 0,763 | no |
| DNAH14 | Irinotecan | 4 | 31 | 0,776 | 0,776 | no |
| HNF1A | Fluorouracil | 6 | 29 | 0,776 | 0,776 | no |
| HNF1A | Oxaliplatin | 4 | 17 | 0,574 | 0,776 | no |
| PABPC3 | Fluorouracil | 19 | 16 | 0,778 | 0,778 | no |
| PABPC3 | Irinotecan | 18 | 17 | 0,298 | 0,778 | no |
| PABPC3 | Oxaliplatin | 11 | 10 | 0,705 | 0,778 | no |
| SAFB2 | Irinotecan | 7 | 28 | 0,523 | 0,7845 | no |
| PRRC2A | Irinotecan | 6 | 29 | 0,526 | 0,789 | no |
| BACE2 | Irinotecan | 13 | 22 | 0,528 | 0,792 | no |
| BACE2 | Oxaliplatin | 5 | 16 | 0,354 | 0,792 | no |
| KIF1A | Fluorouracil | 6 | 29 | 0,264 | 0,792 | no |
| UBR5 | Irinotecan | 5 | 30 | 0,795 | 0,795 | no |
| UBR5 | Oxaliplatin | 4 | 17 | 0,763 | 0,795 | no |
| RIMBP2 | Irinotecan | 8 | 27 | 0,798 | 0,798 | no |
| RIMBP2 | Oxaliplatin | 5 | 16 | 0,548 | 0,798 | no |
| ABCA13 | Irinotecan | 9 | 26 | 0,806 | 0,806 | no |
| COL4A6 | Irinotecan | 6 | 29 | 0,81 | 0,81 | no |
| TENM2 | Fluorouracil | 6 | 29 | 0,81 | 0,81 | no |
| TENM2 | Irinotecan | 6 | 29 | 0,615 | 0,81 | no |
| TENM2 | Oxaliplatin | 2 | 19 | 0,286 | 0,81 | no |
| KIF1A | Irinotecan | 6 | 29 | 0,81 | 0,81 | no |
| KIF1A | Oxaliplatin | 5 | 16 | 0,603 | 0,81 | no |
| PAK2 | Irinotecan | 10 | 25 | 0,812 | 0,812 | no |
| PAK2 | Oxaliplatin | 8 | 13 | 0,697 | 0,812 | no |
| AR | Irinotecan | 5 | 30 | 0,689 | 0,814 | no |
| AR | Oxaliplatin | 3 | 18 | 0,814 | 0,814 | no |
| RAI1 | Fluorouracil | 5 | 30 | 0,689 | 0,814 | no |
| RAI1 | Irinotecan | 5 | 30 | 0,358 | 0,814 | no |
| RAI1 | Oxaliplatin | 3 | 18 | 0,814 | 0,814 | no |
| TGFBR2 | Irinotecan | 5 | 30 | 0,689 | 0,814 | no |
| TGFBR2 | Oxaliplatin | 3 | 18 | 0,814 | 0,814 | no |
| ZNF469 | Irinotecan | 4 | 31 | 0,736 | 0,814 | no |
| ZNF469 | Oxaliplatin | 3 | 18 | 0,814 | 0,814 | no |
| LRP2 | Irinotecan | 5 | 30 | 0,724 | 0,814 | no |
| LRP2 | Oxaliplatin | 3 | 18 | 0,814 | 0,814 | no |
| MSH6 | Irinotecan | 4 | 31 | 0,736 | 0,814 | no |
| MSH6 | Oxaliplatin | 3 | 18 | 0,814 | 0,814 | no |
| ACVR2A | Irinotecan | 5 | 30 | 0,689 | 0,814 | no |
| ACVR2A | Oxaliplatin | 3 | 18 | 0,814 | 0,814 | no |
| ATN1 | Fluorouracil | 7 | 28 | 0,606 | 0,814 | no |
| ATN1 | Oxaliplatin | 3 | 18 | 0,814 | 0,814 | no |
| KIT | Fluorouracil | 4 | 31 | 0,816 | 0,816 | no |
| KIT | Irinotecan | 4 | 31 | 0,312 | 0,816 | no |
| KIT | Oxaliplatin | 2 | 19 | 0,61 | 0,816 | no |
| ABCF1 | Irinotecan | 13 | 22 | 0,824 | 0,824 | no |
| CFAP46 | Irinotecan | 5 | 30 | 0,832 | 0,832 | no |
| ZNF831 | Irinotecan | 5 | 30 | 0,832 | 0,832 | no |
| NASP | Fluorouracil | 6 | 29 | 0,47 | 0,834 | no |
| NASP | Irinotecan | 5 | 30 | 0,556 | 0,834 | no |
| HTT | Oxaliplatin | 5 | 16 | 0,842 | 0,842 | no |
| BAX | Fluorouracil | 7 | 28 | 0,757 | 0,842 | no |
| BAX | Irinotecan | 8 | 27 | 0,829 | 0,842 | no |
| BAX | Oxaliplatin | 5 | 16 | 0,842 | 0,842 | no |
| APOB | Irinotecan | 6 | 29 | 0,844 | 0,844 | no |
| PABPC1 | Fluorouracil | 22 | 13 | 0,282 | 0,846 | no |
| HRNR | Oxaliplatin | 6 | 15 | 0,85 | 0,85 | no |
| KRAS | Irinotecan | 7 | 28 | 0,853 | 0,853 | no |
| UNC80 | Irinotecan | 5 | 30 | 0,289 | 0,867 | no |
| HELZ2 | Fluorouracil | 7 | 28 | 0,578 | 0,867 | no |
| HELZ2 | Oxaliplatin | 5 | 16 | 0,313 | 0,867 | no |
| RP1L1 | Irinotecan | 5 | 30 | 0,869 | 0,869 | no |
| DCHS2 | Fluorouracil | 5 | 30 | 0,869 | 0,869 | no |
| UNC80 | Fluorouracil | 5 | 30 | 0,869 | 0,869 | no |
| UNC80 | Oxaliplatin | 3 | 18 | 0,814 | 0,869 | no |
| ADAMTS20 | Fluorouracil | 6 | 29 | 0,878 | 0,878 | no |
| ADAMTS20 | Oxaliplatin | 4 | 17 | 0,698 | 0,878 | no |
| GDF11 | Fluorouracil | 6 | 29 | 0,878 | 0,878 | no |
| GDF11 | Irinotecan | 7 | 28 | 0,853 | 0,878 | no |
| RBMX | Fluorouracil | 7 | 28 | 0,853 | 0,885 | no |
| RBMX | Irinotecan | 7 | 28 | 0,885 | 0,885 | no |
| RBMX | Oxaliplatin | 4 | 17 | 0,317 | 0,885 | no |
| HELZ2 | Irinotecan | 7 | 28 | 0,885 | 0,885 | no |
| TRIM48 | Fluorouracil | 4 | 31 | 0,776 | 0,887 | no |
| TRIM48 | Irinotecan | 6 | 29 | 0,37 | 0,887 | no |
| TRIM48 | Oxaliplatin | 3 | 18 | 0,887 | 0,887 | no |
| RNF213 | Fluorouracil | 8 | 27 | 0,891 | 0,891 | no |
| SACS | Irinotecan | 5 | 30 | 0,869 | 0,897 | no |
| SACS | Oxaliplatin | 4 | 17 | 0,897 | 0,897 | no |
| CACNA1A | Fluorouracil | 6 | 29 | 0,526 | 0,905 | no |
| CACNA1A | Irinotecan | 5 | 30 | 0,759 | 0,905 | no |
| CACNA1A | Oxaliplatin | 5 | 16 | 0,905 | 0,905 | no |
| NASP | Oxaliplatin | 5 | 16 | 0,905 | 0,905 | no |
| MT-CYB | Irinotecan | 7 | 28 | 0,606 | 0,909 | no |
| MT-CYB | Oxaliplatin | 3 | 18 | 0,471 | 0,909 | no |
| IGLV5-45 | Oxaliplatin | 6 | 15 | 0,91 | 0,91 | no |
| SALL1 | Fluorouracil | 6 | 29 | 0,913 | 0,913 | no |
| SALL1 | Irinotecan | 6 | 29 | 0,678 | 0,913 | no |
| SALL1 | Oxaliplatin | 4 | 17 | 0,517 | 0,913 | no |
| RBMXL1 | Fluorouracil | 6 | 29 | 0,913 | 0,913 | no |
| MT-CYB | Fluorouracil | 7 | 28 | 0,918 | 0,918 | no |
| FRMPD4 | Fluorouracil | 6 | 29 | 0,615 | 0,9225 | no |
| ZFHX4 | Irinotecan | 3 | 32 | 0,93 | 0,93 | no |
| ZFHX4 | Oxaliplatin | 3 | 18 | 0,814 | 0,93 | no |
| BACE2 | Fluorouracil | 13 | 22 | 0,932 | 0,932 | no |
| ERBB2 | Fluorouracil | 4 | 31 | 0,938 | 0,938 | no |
| H1-8 | Fluorouracil | 5 | 30 | 0,944 | 0,944 | no |
| PIK3CA | Irinotecan | 6 | 29 | 0,948 | 0,948 | no |
| PIK3CA | Oxaliplatin | 6 | 15 | 0,91 | 0,948 | no |
| TNS1 | Fluorouracil | 6 | 29 | 0,948 | 0,948 | no |
| TNS1 | Irinotecan | 6 | 29 | 0,776 | 0,948 | no |
| TNS1 | Oxaliplatin | 4 | 17 | 0,897 | 0,948 | no |
| BLTP1 | Irinotecan | 6 | 29 | 0,948 | 0,948 | no |
| BLTP1 | Oxaliplatin | 4 | 17 | 0,897 | 0,948 | no |
| PRRC2A | Fluorouracil | 6 | 29 | 0,948 | 0,948 | no |
| PTEN | Oxaliplatin | 2 | 19 | 0,952 | 0,952 | no |
| PABPC1 | Irinotecan | 22 | 13 | 0,959 | 0,959 | no |
| PABPC1 | Oxaliplatin | 12 | 9 | 0,702 | 0,959 | no |
| ALK | Fluorouracil | 1 | 34 | 0,961 | 0,961 | no |
| ALK | Irinotecan | 2 | 33 | 0,594 | 0,961 | no |
| ALK | Oxaliplatin | 1 | 20 | 0,762 | 0,961 | no |
| FRMPD4 | Oxaliplatin | 3 | 18 | 0,962 | 0,962 | no |
| TCF7L2 | Oxaliplatin | 4 | 17 | 0,965 | 0,965 | no |
| MUC19 | Fluorouracil | 15 | 20 | 0,855 | 0,971 | no |
| MUC19 | Oxaliplatin | 7 | 14 | 0,971 | 0,971 | no |
| ZFHX3 | Irinotecan | 6 | 29 | 0,983 | 0,983 | no |
| ZFHX3 | Oxaliplatin | 5 | 16 | 0,719 | 0,983 | no |
| KMT2C | Fluorouracil | 8 | 27 | 0,768 | 0,984 | no |
| KMT2C | Irinotecan | 7 | 28 | 0,984 | 0,984 | no |
| KMT2C | Oxaliplatin | 6 | 15 | 0,91 | 0,984 | no |
| HLA-DRB5 | Oxaliplatin | 5 | 16 | 1 | 1 | no |
| SAFB2 | Oxaliplatin | 5 | 16 | 1 | 1 | no |
| CSMD1 | Irinotecan | 7 | 28 | 0,885 | 1 | no |
| CSMD1 | Oxaliplatin | 5 | 16 | 1 | 1 | no |
| FREM2 | Oxaliplatin | 3 | 18 | 1 | 1 | no |
| CASP5 | Oxaliplatin | 4 | 17 | 1 | 1 | no |
| MYO15B | Fluorouracil | 4 | 31 | 1 | 1 | no |
| KCTD8 | Fluorouracil | 6 | 29 | 1 | 1 | no |
| KCTD8 | Irinotecan | 6 | 29 | 0,526 | 1 | no |
| KCTD8 | Oxaliplatin | 1 | 20 | 0,667 | 1 | no |
| BRAF | Fluorouracil | 5 | 30 | 0,724 | 1 | no |
| BRAF | Irinotecan | 5 | 30 | 1 | 1 | no |
| BRAF | Oxaliplatin | 4 | 17 | 0,897 | 1 | no |
| HSPD1 | Irinotecan | 11 | 24 | 0,901 | 1 | no |
| HSPD1 | Oxaliplatin | 8 | 13 | 1 | 1 | no |

**Supplementary Table 2.** Molecular classification of PDTOs.

| pdto_id | prediction | d.iCMS2 | d.iCMS3 | p.value | FDR |
| --- | --- | --- | --- | --- | --- |
| CRC561ORG | iCMS2 | 0,586752641 | 0,809766225 | 0,001 | 0,001347826 |
| CRC997ORG | iCMS2 | 0,679486994 | 0,733687552 | 0,037962038 | 0,046149929 |
| CRC321ORG | iCMS2 | 0,644786623 | 0,764362617 | 0,001 | 0,001347826 |
| CRC153ORG | iCMS2 | 0,675036576 | 0,737784265 | 0,016983017 | 0,021488715 |
| CRC074ORG | iCMS3 | 0,889362569 | 0,457202603 | 0,001 | 0,001347826 |
| CRC228ORG | iCMS2 | 0,597122519 | 0,802150046 | 0,001 | 0,001347826 |
| CRC146ORG | iCMS3 | 0,925504146 | 0,378737475 | 0,001 | 0,001347826 |
| CRC634ORG | iCMS2 | 0,548570159 | 0,836104528 | 0,001 | 0,001347826 |
| CRC049ORG | iCMS2 | 0,571346785 | 0,820708749 | 0,001 | 0,001347826 |
| CRC128ORG | iCMS2 | 0,624654539 | 0,780901215 | 0,001 | 0,001347826 |
| CRC303ORG | iCMS3 | 0,910194859 | 0,414180296 | 0,001 | 0,001347826 |
| CRC024ORG | iCMS2 | 0,627117856 | 0,778924383 | 0,001 | 0,001347826 |
| CRC839ORG | iCMS2 | 0,650662263 | 0,759367249 | 0,001 | 0,001347826 |
| CRC356ORG_1 | Undefined | 0,716020953 | 0,698078789 | 0,512487512 | 0,557442557 |
| CRC165ORG | iCMS2 | 0,565828623 | 0,824522874 | 0,001 | 0,001347826 |
| CRC622ORG | iCMS2 | 0,668955256 | 0,743302674 | 0,011988012 | 0,015484515 |
| CRC882ORG | iCMS2 | 0,616580158 | 0,787292137 | 0,001 | 0,001347826 |
| CRC879ORG | iCMS3 | 0,756683605 | 0,653781249 | 0,001 | 0,001347826 |
| CRC899ORG | iCMS3 | 0,861140663 | 0,508366756 | 0,001 | 0,001347826 |
| CRC297ORG | iCMS3 | 0,840796836 | 0,541350792 | 0,001 | 0,001347826 |
| CRC986ORG | iCMS2 | 0,589385332 | 0,807852048 | 0,001 | 0,001347826 |
| CRC621ORG | iCMS2 | 0,554704112 | 0,832047684 | 0,001 | 0,001347826 |
| CRC212ORG | iCMS2 | 0,667841693 | 0,744303347 | 0,004995005 | 0,006589156 |
| CRC930ORG_2 | Undefined | 0,712781788 | 0,701385859 | 0,669330669 | 0,715491405 |
| CRC860ORG | Undefined | 0,709013446 | 0,705194962 | 0,884115884 | 0,913586414 |
| CRC259ORG | iCMS3 | 0,914636273 | 0,404277737 | 0,001 | 0,001347826 |
| CRC314ORG | iCMS2 | 0,637794813 | 0,770206321 | 0,001 | 0,001347826 |
| CRC481ORG | iCMS2 | 0,595906124 | 0,803054102 | 0,001 | 0,001347826 |
| CRC298ORG | Undefined | 0,726594794 | 0,68706623 | 0,134865135 | 0,152029788 |
| CRC989ORG | iCMS2 | 0,556983618 | 0,830523479 | 0,001 | 0,001347826 |
| CRC158ORG | iCMS2 | 0,579179311 | 0,815200175 | 0,001 | 0,001347826 |
| CRC299ORG | iCMS2 | 0,599392069 | 0,800455587 | 0,001 | 0,001347826 |
| CRC911ORG | Undefined | 0,730992158 | 0,682385862 | 0,067932068 | 0,079467702 |
| CRC967ORG | iCMS2 | 0,595506728 | 0,80335032 | 0,001 | 0,001347826 |
| CRC648ORG | iCMS2 | 0,603906508 | 0,797055161 | 0,001 | 0,001347826 |
| CRC836ORG | Undefined | 0,707908239 | 0,706304414 | 0,952047952 | 0,952047952 |
| CRC994ORG | iCMS3 | 0,892246531 | 0,451548588 | 0,001 | 0,001347826 |
| CRC197ORG | iCMS2 | 0,598363809 | 0,801224533 | 0,001 | 0,001347826 |
| CRC516ORG | iCMS3 | 0,790055341 | 0,613035527 | 0,001 | 0,001347826 |
| CRC226ORG | iCMS2 | 0,534188295 | 0,845365522 | 0,001 | 0,001347826 |
| CRC504ORG | Undefined | 0,728810218 | 0,684715755 | 0,07992008 | 0,091760092 |
| CRC946ORG | iCMS3 | 0,757704839 | 0,652597408 | 0,001 | 0,001347826 |
| CRC262ORG | iCMS2 | 0,611458384 | 0,791276592 | 0,001 | 0,001347826 |
| CRC390ORG | Undefined | 0,701382935 | 0,712784665 | 0,682317682 | 0,717011802 |
| CRC372ORG | iCMS2 | 0,542036087 | 0,840355211 | 0,001 | 0,001347826 |
| CRC003ORG | iCMS2 | 0,561671672 | 0,827360219 | 0,001 | 0,001347826 |
| CRC374ORG | iCMS2 | 0,650700382 | 0,759334586 | 0,001 | 0,001347826 |
| CRC405ORG | iCMS2 | 0,582897719 | 0,812545537 | 0,001 | 0,001347826 |
| CRC770ORG | iCMS2 | 0,519951427 | 0,854195829 | 0,001 | 0,001347826 |
| CRC870ORG | iCMS3 | 0,916027667 | 0,401115088 | 0,001 | 0,001347826 |
| CRC698ORG | iCMS2 | 0,633443001 | 0,773789354 | 0,001 | 0,001347826 |
| CRC650ORG | iCMS2 | 0,590270705 | 0,807205361 | 0,001 | 0,001347826 |
| CRC040ORG | iCMS3 | 0,895146099 | 0,445772882 | 0,001 | 0,001347826 |
| CRC973ORG | iCMS3 | 0,859614923 | 0,510942448 | 0,001 | 0,001347826 |
| CRC033ORG | iCMS3 | 0,787915876 | 0,615782894 | 0,001 | 0,001347826 |
| CRC103ORG_2 | Undefined | 0,705782086 | 0,708429 | 0,922077922 | 0,937193954 |
| CRC109ORG | Undefined | 0,724456642 | 0,689320371 | 0,188811189 | 0,209040959 |
| CRC669ORG | iCMS2 | 0,625986263 | 0,779834084 | 0,001 | 0,001347826 |
| CRC076ORG | iCMS3 | 0,778362795 | 0,627814749 | 0,001 | 0,001347826 |
| CRC777ORG | iCMS2 | 0,676208137 | 0,736710632 | 0,032967033 | 0,040879121 |
| CRC491ORG | iCMS2 | 0,605332286 | 0,795972879 | 0,001 | 0,001347826 |
| CRC357ORG | Undefined | 0,68108556 | 0,732203838 | 0,057942058 | 0,069084761 |

**Supplementary Table 3.**
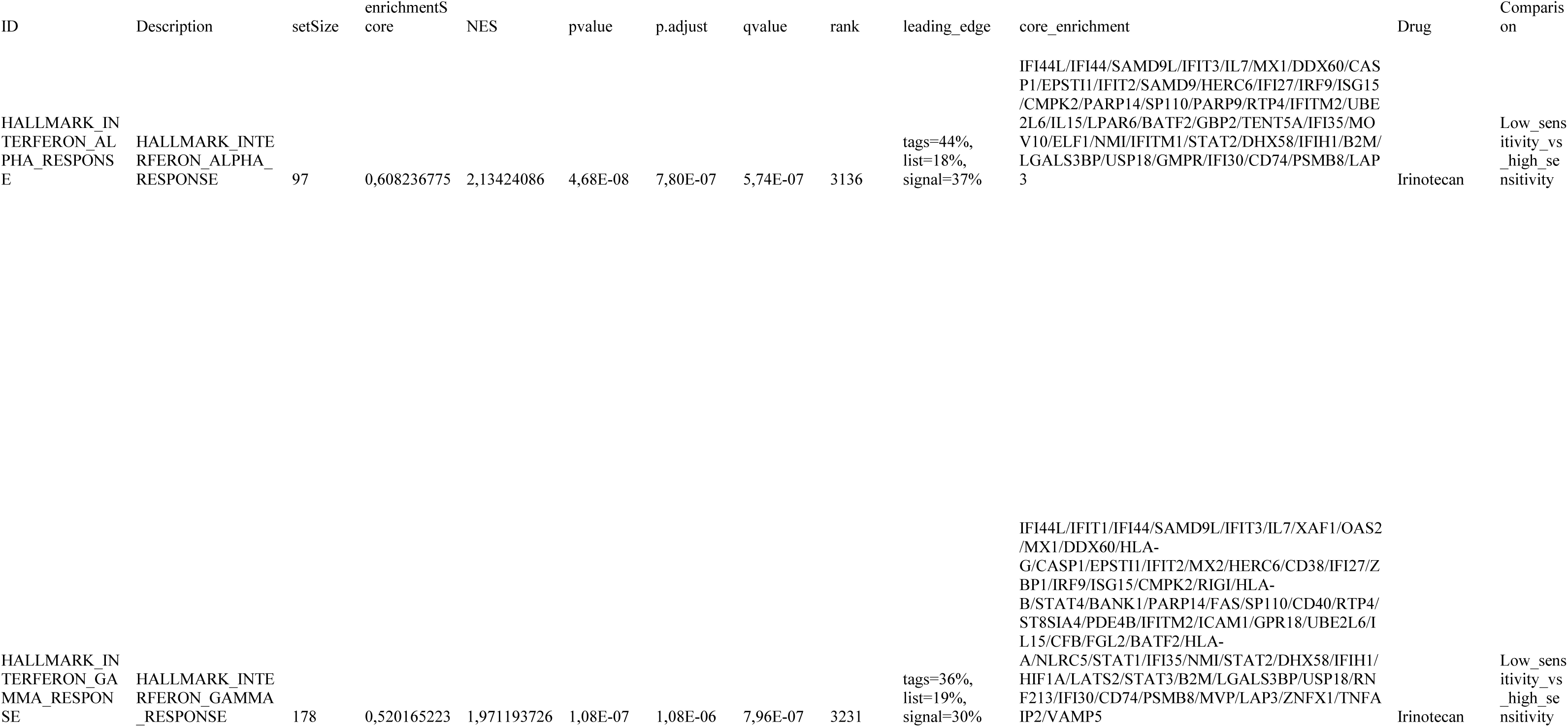
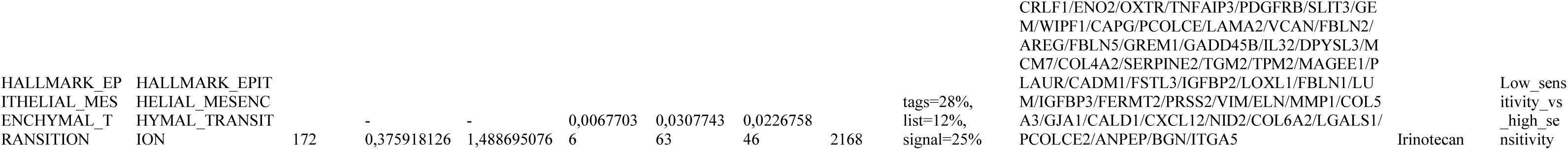
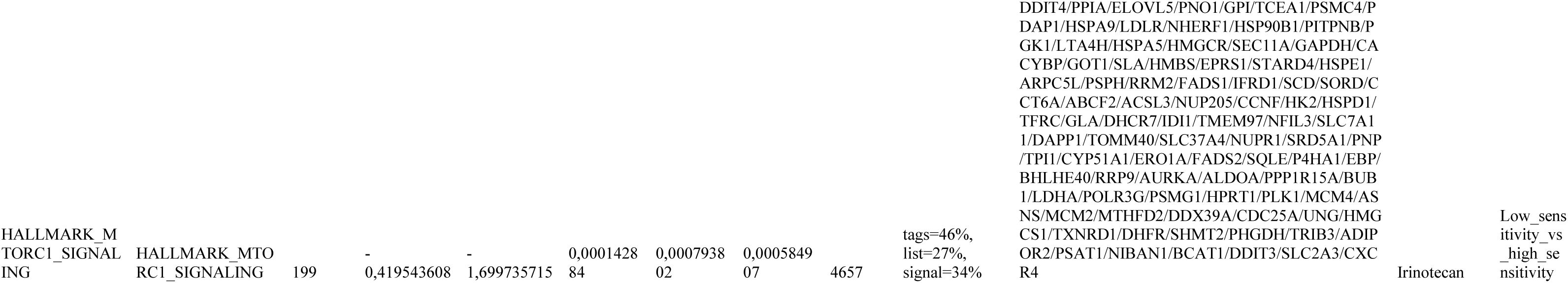
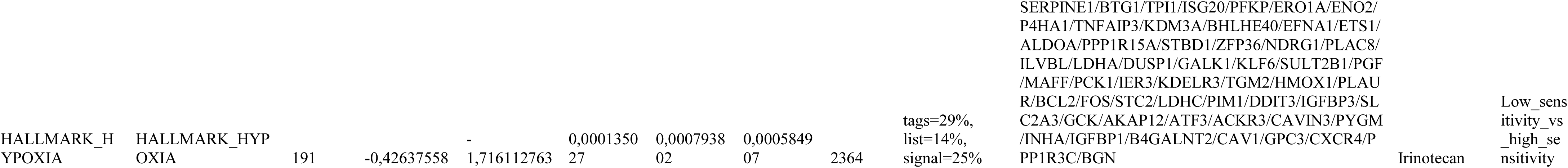
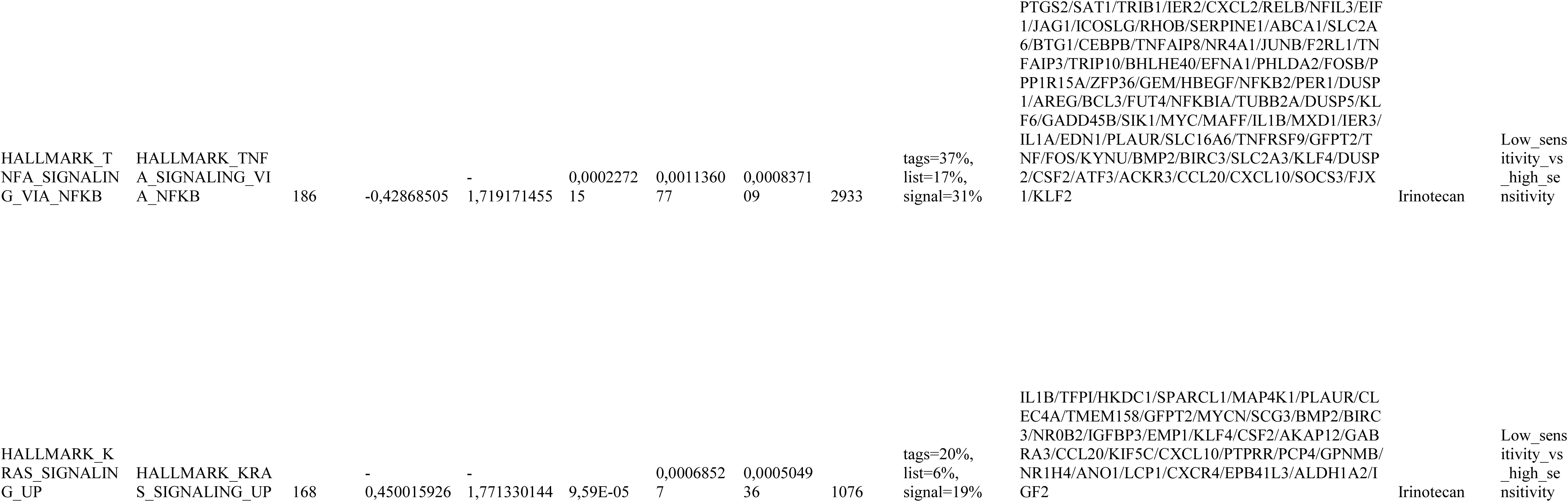
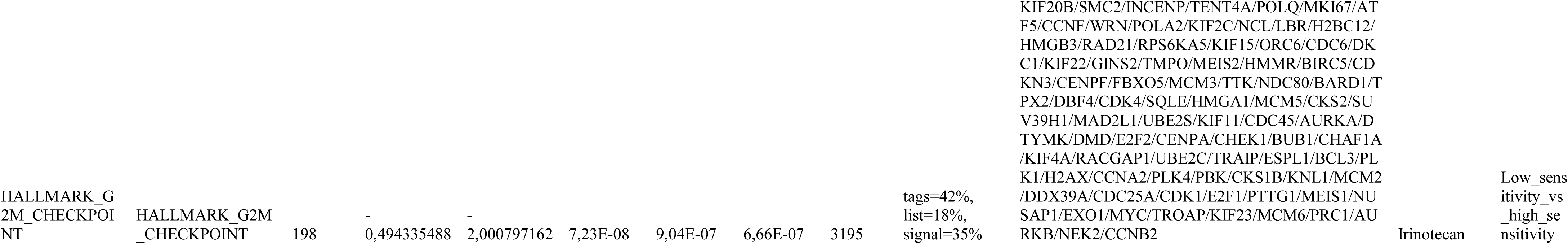
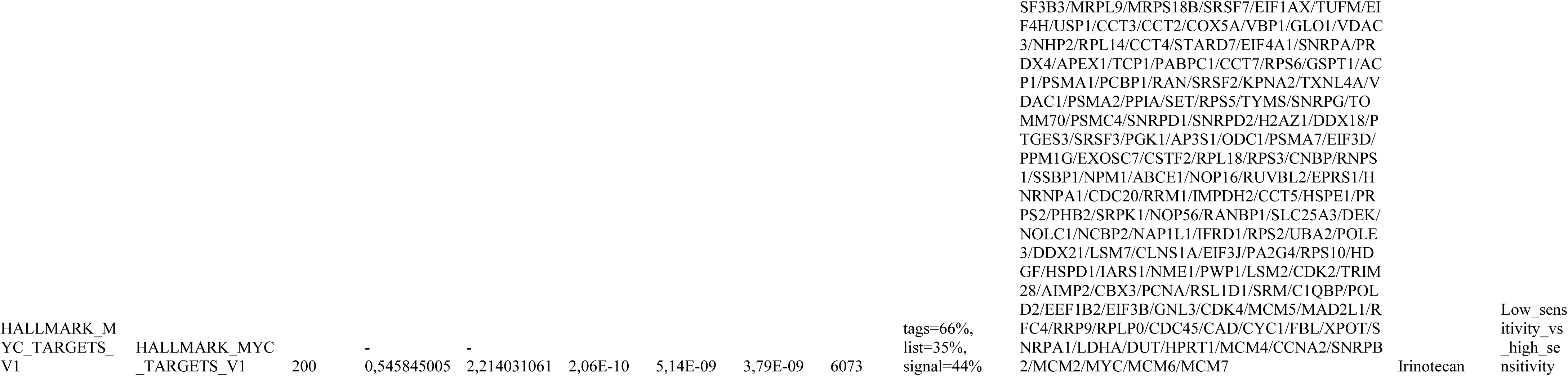
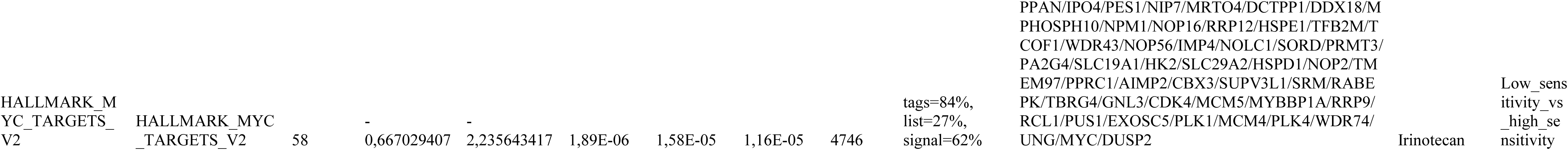
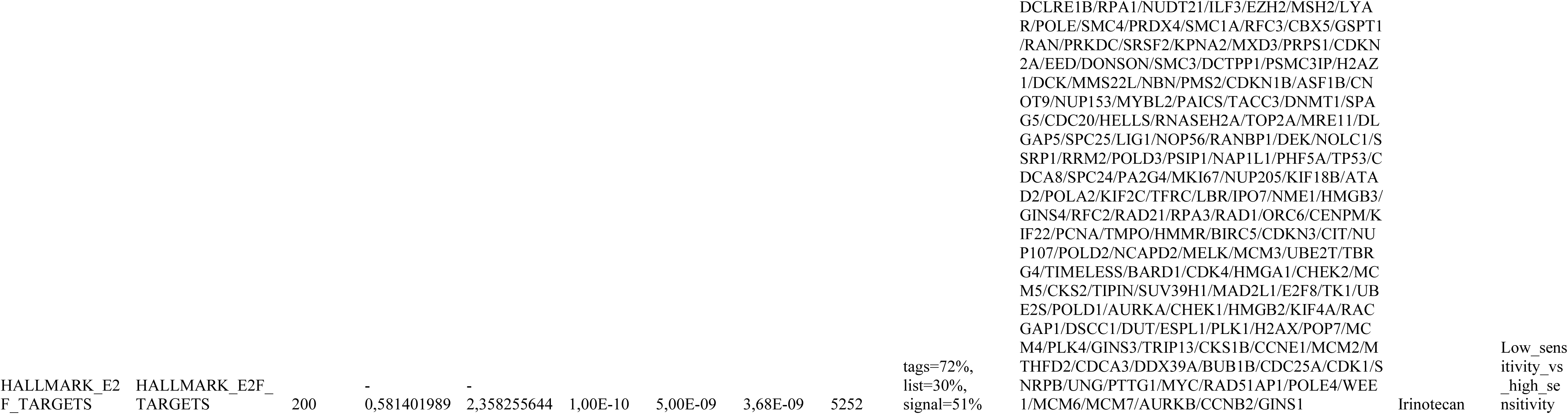
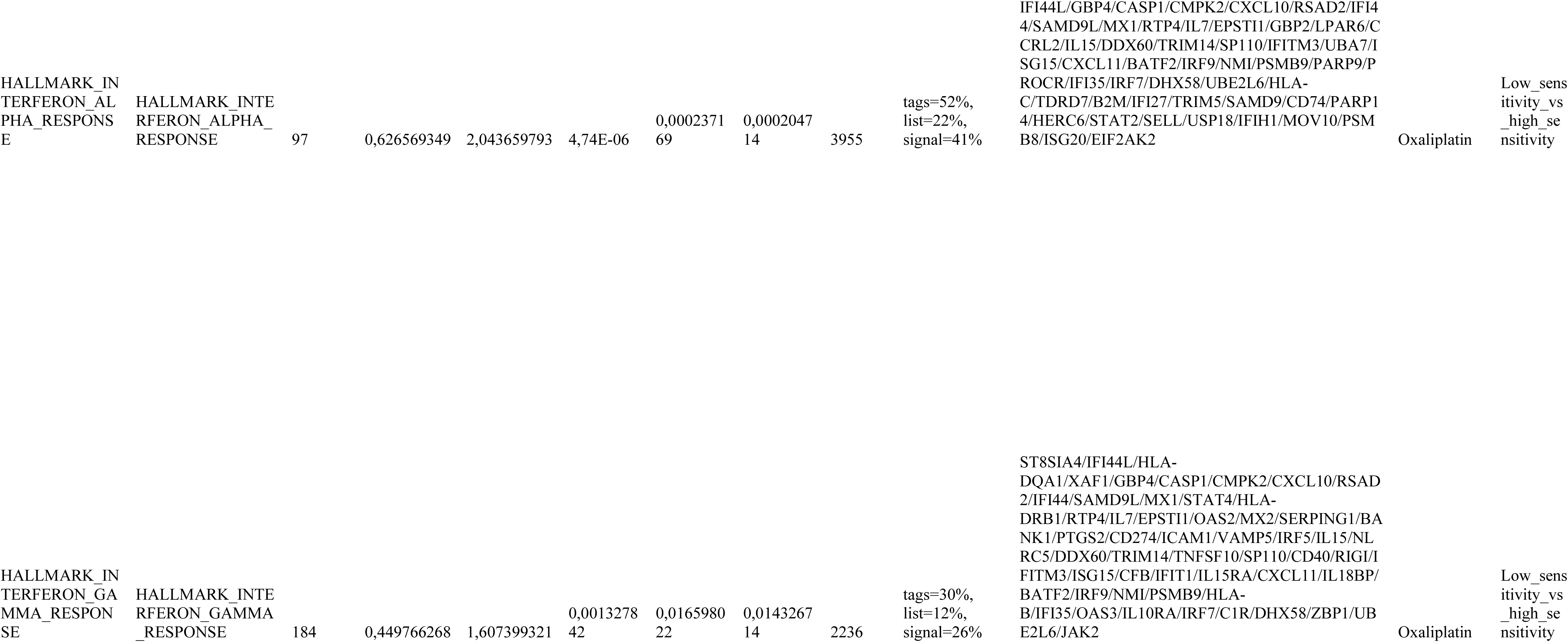
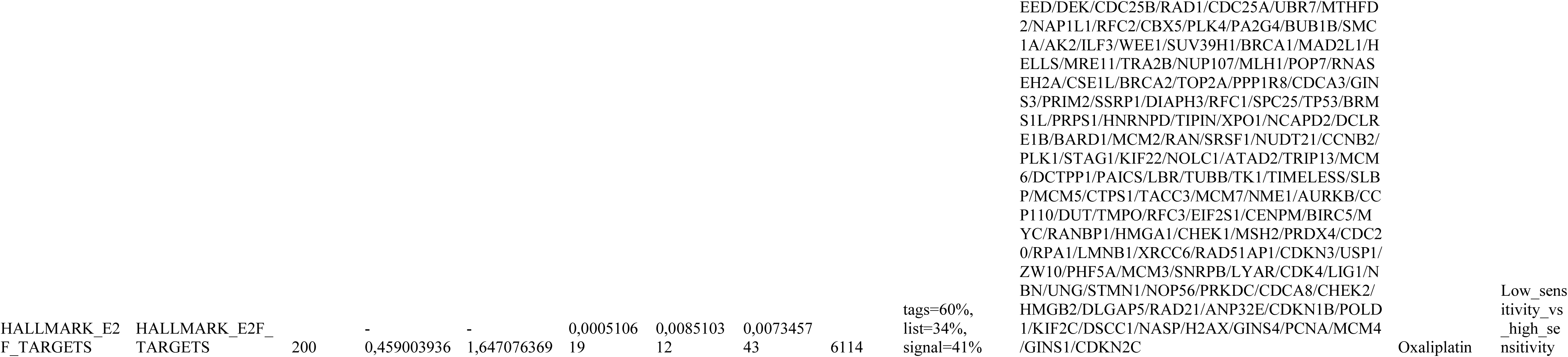
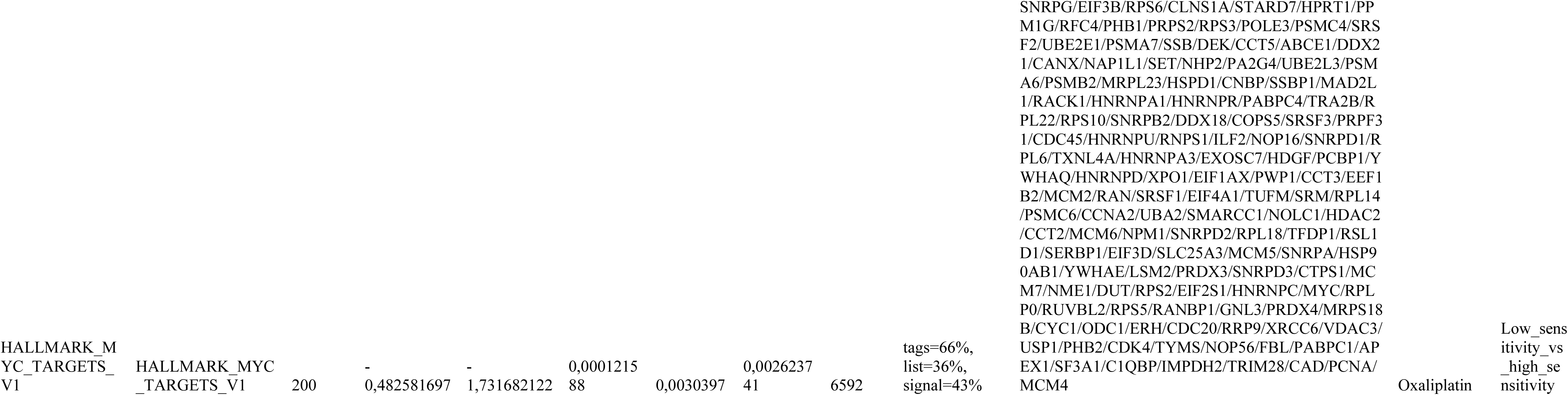
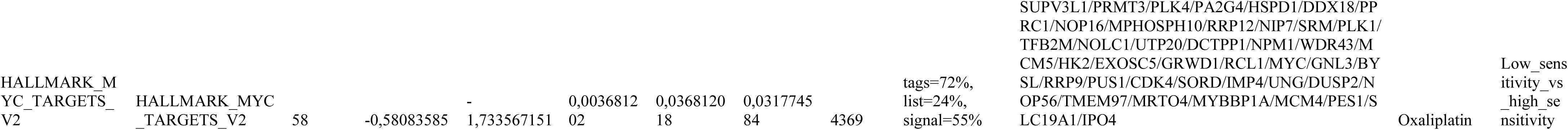
Aggregated GSEA results.

## REFERENCES

1. Wang, R. et al. Systematic evaluation of colorectal cancer organoid system by single-cell RNA-Seq analysis. Genome Biol. 23, 106 (2022).

2. Fujii, M. et al. A Colorectal Tumor Organoid Library Demonstrates Progressive Loss of Niche Factor Requirements during Tumorigenesis. Cell Stem Cell 18, 827–838 (2016).

3. Mao, Y. et al. Drug repurposing screening and mechanism analysis based on human colorectal cancer organoids. Protein Cell 15, 285–304 (2024).

4. van de Wetering, M. et al. Prospective Derivation of a Living Organoid Biobank of Colorectal Cancer Patients. Cell 161, 933–945 (2015).

5. Roper, J. et al. In vivo genome editing and organoid transplantation models of colorectal cancer and metastasis. Nat. Biotechnol. 35, 569–576 (2017).

6. Ooft, S. N. et al. Patient-derived organoids can predict response to chemotherapy in metastatic colorectal cancer patients. Sci. Transl. Med. 11, eaay2574 (2019).

7. Yao, Y. et al. Patient-Derived Organoids Predict Chemoradiation Responses of Locally Advanced Rectal Cancer. Cell Stem Cell 26, 17–26.e6 (2020).

8. Chalabi, M. et al. Neoadjuvant immunotherapy leads to pathological responses in MMR-proficient and MMR-deficient early-stage colon cancers. Nat. Med. 26, 566–576 (2020).

9. Hao, M. et al. Patient-Derived Organoid Model in the Prediction of Chemotherapeutic Drug Response in Colorectal Cancer. ACS Biomater. Sci. Eng. 8, 3515–3525 (2022).

10. Mo, S. et al. Patient-Derived Organoids from Colorectal Cancer with Paired Liver Metastasis Reveal Tumor Heterogeneity and Predict Response to Chemotherapy. Adv. Sci. 9, 2204097 (2022).

11. Amiryousefi, A., Williams, B., Jafari, M. & Tang, J. The ENDS of assumptions: an online tool for the epistemic non-parametric drug–response scoring. Bioinformatics 38, 3132–3133 (2022).

12. Mpindi, J. P. et al. Consistency in drug response profiling. Nature 540, E5–E6 (2016).

13. Brooks, E. A. et al. Applicability of drug response metrics for cancer studies using biomaterials. Philos. Trans. R. Soc. B Biol. Sci. 374, 20180226 (2019).

14. Love, M. I., Huber, W. & Anders, S. Moderated estimation of fold change and dispersion for RNA-seq data with DESeq2. Genome Biol. 15, 550 (2014).

15. Durinck, S., Spellman, P. T., Birney, E. & Huber, W. Mapping identifiers for the integration of genomic datasets with the R/Bioconductor package biomaRt. Nat. Protoc. 4, 1184–1191 (2009).

16. Blighe, K. EnhancedVolcano. Bioconductor 10.18129/B9.BIOC.ENHANCEDVOLCANO (2018).

17. Badia-i-Mompel, P., et al. decoupleR: ensemble of computational methods to infer biological activities from omics data. *Bioinforma. Adv*. 2, vbac016 (2022).

18. Schubert, M. et al. Perturbation-response genes reveal signaling footprints in cancer gene expression. Nat. Commun. 9, 20 (2018).

19. Korotkevich, G. et al. Fast gene set enrichment analysis. Preprint at 10.1101/060012 (2016).

20. Yu, G., Wang, L.-G., Han, Y. & He, Q.-Y. clusterProfiler: an R Package for Comparing Biological Themes Among Gene Clusters. OMICS J. Integr. Biol. 16, 284–287 (2012).

21. Liberzon, A. et al. The Molecular Signatures Database Hallmark Gene Set Collection. Cell Syst. 1, 417–425 (2015).

22. Eide, P. W., Bruun, J., Lothe, R. A. & Sveen, A. CMScaller: an R package for consensus molecular subtyping of colorectal cancer pre-clinical models. Sci. Rep. 7, 16618 (2017).

23. Joanito, I. et al. Single-cell and bulk transcriptome sequencing identifies two epithelial tumor cell states and refines the consensus molecular classification of colorectal cancer. Nat. Genet. 54, 963–975 (2022).

24. Qin, P. et al. Cancer-associated fibroblasts undergoing neoadjuvant chemotherapy suppress rectal cancer revealed by single-cell and spatial transcriptomics. Cell Rep. Med. 4, 101231 (2023).

25. Domínguez Conde, C., et al. Cross-tissue immune cell analysis reveals tissue-specific features in humans. Science 376, eabl5197 (2022).

26. Guinney, J. et al. The consensus molecular subtypes of colorectal cancer. Nat. Med. 21, 1350–1356 (2015).

27. Yang, W. et al. Genomics of Drug Sensitivity in Cancer (GDSC): a resource for therapeutic biomarker discovery in cancer cells. Nucleic Acids Res. 41, D955–D961 (2012).

28. Iorio, F. et al. A Landscape of Pharmacogenomic Interactions in Cancer. Cell 166, 740– 754 (2016).

29. De Neergaard, M. et al. Epithelial-Stromal Interaction 1 (EPSTI1) Substitutes for Peritumoral Fibroblasts in the Tumor Microenvironment. Am. J. Pathol. 176, 1229–1240 (2010).

30. Rao, C. et al. KSR1- and ERK-dependent translational regulation of the epithelial-to-mesenchymal transition. eLife 10, e66608 (2021).

31. Seppälä, T. T. et al. Patient-derived Organoid Pharmacotyping is a Clinically Tractable Strategy for Precision Medicine in Pancreatic Cancer. Ann. Surg. 272, 427–435 (2020).

32. Musella, M. et al. Type I IFNs promote cancer cell stemness by triggering the epigenetic regulator KDM1B. Nat. Immunol. 23, 1379–1392 (2022).

33. Yamashita, N. et al. Abstract PO3-13-07: MUC1-C INTEGRATES CHRONIC ACTIVATION OF INTERFERON PATHWAYS WITH CHROMATIN REMODELING IN TREATMENT RESISTANCE OF TRIPLE-NEGATIVE BREAST CANCER. Cancer Res. 84, PO3–13-07-PO3-13–07 (2024).

34. Ganesh, K. et al. A rectal cancer organoid platform to study individual responses to chemoradiation. Nat. Med. 25, 1607–1614 (2019).

35. Janakiraman, H. et al. Modeling rectal cancer to advance neoadjuvant precision therapy. Int. J. Cancer 147, 1405–1418 (2020).

36. Li, N. et al. Mapping and modeling human colorectal carcinoma interactions with the tumor microenvironment. Nat. Commun. 14, 7915 (2023).

